# Persistent and recyclable myosin II minifilaments drive cortical contractility in the early *C. elegans* embryo

**DOI:** 10.64898/2026.11.04.764377

**Authors:** Karen Lorena Chauca Espinoza, Shashi Kumar Suman, Teresa Ferraro, Camille Plancke, Pierre Mahou, Emmanuel Beaurepaire, Tae Yoon Kim, François Robin

**Author notes:** Equal contribution (Karen Lorena Chauca Espinoza, Shashi Kumar Suman).

## Abstract

Non-muscle myosin II is a key regulator of cell and tissue morphogenesis, yet the cortical dynamics of myosin minifilaments *in vivo* remain poorly understood. Here, we combine SIM-TIRF and single-molecule TIRF microscopy to characterize the structure and dynamics of myosin minifilaments in an early developing embryo. High-temporal-resolution SIM-TIRF imaging revealed diverse minifilament behaviors, including partition-like events, long-range alignment of multiple minifilaments into collective structures we called proto-stress fibers, partial unbinding, and sequential unbinding and rebinding to the cortex. Photoconversion experiments showed that minifilaments persist as intact units after cortical unbinding, diffuse through the cytoplasm, and can be recycled over long distances rather than disassembling upon detachment. Despite these dynamic behaviors, minifilaments are predominantly stationary when bound to actin. Numerical simulations demonstrated that this static behavior is essential to sustain cortical stress within a continuously remodeling actin network. Using myosin motor-domain mutants, we further investigated the contribution of motor activity to these behaviors. Together, our results reveal that myosin minifilaments can persist as stable units while cycling between the cortex and cytoplasm, and that their predominantly static behavior when bound to actin enables sustained force generation in a continuously renewing cortical network. These findings provide new insights into myosin organization, recycling, and force generation during early embryonic development.

## INTRODUCTION

Cell shape changes, cell division, and migration rely on the precise spatial and temporal control of contractile forces generated by the actomyosin cortex, a thin and dynamic network underlying the plasma membrane. At the core of this system, non-muscle myosin II converts chemical energy into mechanical work by interacting with F-actin, thereby driving cortical tension and cytoskeletal remodeling. While the molecular composition and biochemical regulation of myosin II have been extensively characterized, how the minimal contractile unit, the myosin minifilament, is organized and behaves in living cells remains poorly understood. A central unresolved question is whether cortical myosin II operates through dynamic cycles of minifilament assembly and disassembly, or through preassembled structural units that persist and are reused over time.

Classical models largely assume that the bipolar filament assembly is a process that emerges immediately after the phosphorylation-mediated activation of the Regulatory Light Chain (RLC) (Adelstein & Anne Conti, 1975; Craig et al., 1983; Kendrick-Jones et al., 1987) downstream of pathways such as RhoA–ROCK (Amano et al., 1996; Totsukawa et al., 2000), implying that contractility emerges from continuous turnover of myosin structures. More recent study has refined this view by showing that filament assembly is not solely dictated by phosphorylation-dependent activation, but also by local increases in cytosolic myosin concentration and actin-driven spatial confinement, which together promote the formation and amplification of nascent filaments (Quintanilla et al., 2023). Importantly, in *Caenorhabditis elegans*, a cortex-scale model further proposed that myosin filament assembly and disassembly are spatially regulated by RhoA pathway components: the kinase LET-502 (ROCK) promotes myosin activation and minifilament assembly, whereas the myosin phosphatase MEL-11 favors filament disassembly and motor release into a cytoplasmic pool that can be redistributed to sites of contractility (Najafabadi et al., 2022).

Together, these studies support a model in which myosin filament assembly, growth, and turnover are dynamically regulated in space and time. However, direct *in vivo* evidence remains limited, and it is unclear how the structural state of myosin relates to its behavior within the complex cortical environment. Addressing this question requires a detailed understanding of the organization and dynamics of the minimal contractile unit.

At the molecular level, NM2 assembles into bipolar minifilaments composed of ~60 myosin motor domains (Niederman & Pollard, 1975; Billington et al., 2013; Melli et al., 2018) arranged around a central bare zone and oppositely oriented motor domains capable of engaging actin filaments in multiple configurations (Niederman & Pollard, 1975; Billington et al., 2013). *In vitro* reconstitution experiments have shown that minifilaments can move processively along actin depending on motor number (Nagy et al., 2013; Melli et al., 2018). However, these studies have been performed in simplified and highly ordered systems, leaving open how such assemblies behave *in vivo*, where actin networks are disordered, dynamic, and spatially heterogeneous.

Beyond individual minifilaments, NMII contractile systems are organized through collective interactions between multiple bipolar assemblies. At larger spatial scales, minifilaments align along actin filaments to generate contractile structures such as stress fibers, contractile arcs, and cortical actomyosin bundles, enabling force production across extended regions of the cell (Hotulainen & Lappalainen, 2006; Burnette et al., 2014; Lehtimäki et al., 2021; Vitriol et al., 2023). At smaller scales, neighboring minifilaments can undergo lateral association through mechanisms such as concatenation, generating mechanically coupled assemblies of multiple pre-existing contractile units (Shutova et al., 2014; Fenix et al., 2016; Beach et al., 2017). Thus, NMII organization can be regulated not only through changes in individual minifilaments, but also through their collective arrangement and spatial redistribution within the cortical network.

Distinguishing between these possibilities is critical because they imply fundamentally different mechanisms for regulating contractility. If remodeling primarily depends on minifilament assembly and disassembly, force generation would be controlled through continuous production and turnover of the basic contractile unit. Alternatively, if minifilaments persist as stable structures, contractility could instead be regulated through changes in cortical engagement, spatial redistribution, and organization into higher-order assemblies. Despite the importance of this distinction, direct visualization of individual myosin minifilaments *in vivo* has remained technically challenging, leaving unresolved how their structure, persistence, and collective organization contribute to cortical force generation.

Here, we combine TIRF and super-resolution SIM-TIRF microscopy with photoconversion-based pulse–chase analysis in the early *Caenorhabditis elegans* embryo to directly resolve the structure, stoichiometry, and dynamics of individual cortical NMY-2 minifilaments *in vivo*. We show that NM2 forms stereotypical bipolar minifilaments with defined size and composition that persist over time and do not undergo extensive disassembly upon cortical detachment. Instead, these structures behave as stable, reusable units that dynamically engage with the actin cortex. By tracking individual minifilaments, we uncover a diverse repertoire of individual and collective behaviors.

Moreover, taking advantage of the detailed measurements of minifilament kinetics, we used numerical simulations to study the implications of the observed behavior in terms of network stress. Indeed, the prevalence of stationary behavior during actin-binding phases implies a stress magnitude close to its maximal potential.

Together, our findings challenge current models of non-muscle myosin II dynamics in *vivo*. Instead, our results raise the question of how force generation is regulated through the dynamic engagement with actin, spatial organization, and reuse of preassembled myosin minifilaments, providing a new framework for understanding how contractility emerges in living cells.

## RESULTS

### 1. SIM-TIRF super-resolution reveals the nanoscale organization of myosin minifilaments

To observe the structural organization of myosin in the early embryo, we used CRISPR-based genome editing techniques to generate GFP fusions of *mlc-4* (myosin regulatory light chain, MLC-4::eGFP) and *mlc-5* (myosin essential light chain, eGFP::MLC-5) at the endogenous genomic locus **(see Table S1)**, as well as existing fusion strains of myosin heavy chain labelled either in C-terminus or N-terminus (NMY2::GFP and mNG::NMY-2, (Dickinson et al., 2013, 2015; Heppert et al., 2016).

We then combined these strains with an actin marker, LifeAct::mKate2, (Reymann et al., 2016) and visualized myosin and actin localization in the zygote during polarity maintenance, and at the 2-cell stage during interphase and metaphase, using TIRF microscopy. All strains were healthy and displayed a characteristic and stereotypical pattern as previously described **(Fig. S1, Video 1-5)** (Munro et al., 2004; Kumfer et al., 2010; Reymann et al., 2016; Michaux et al., 2018). As described in previous studies, we observed that cortical myosin in the strain expressing NMY-2::GFP, labeled at the core of the myosin minifilaments, resembled “*dots*” bound to the cortical actin network **(Fig. 1A)**. In striking contrast, in all strains labeled at the extremities of the myosin minifilaments (mNG::NMY-2, MLC-4::eGFP and eGFP::MLC-5), isolated minifilaments displayed a “*rod-lik*e” pattern. This pattern enabled direct assessment of minifilament orientation relative to F-actin and discrimination between mono-polar binding and bipolar engagement with the actin cortex **(Fig. 1B-D)**. Taken together, these observations suggest that myosin light and heavy chains, MLC-4, MLC-5 and NMY-2 assemble at the cortex into myosin minifilaments.

**Figure 1.**
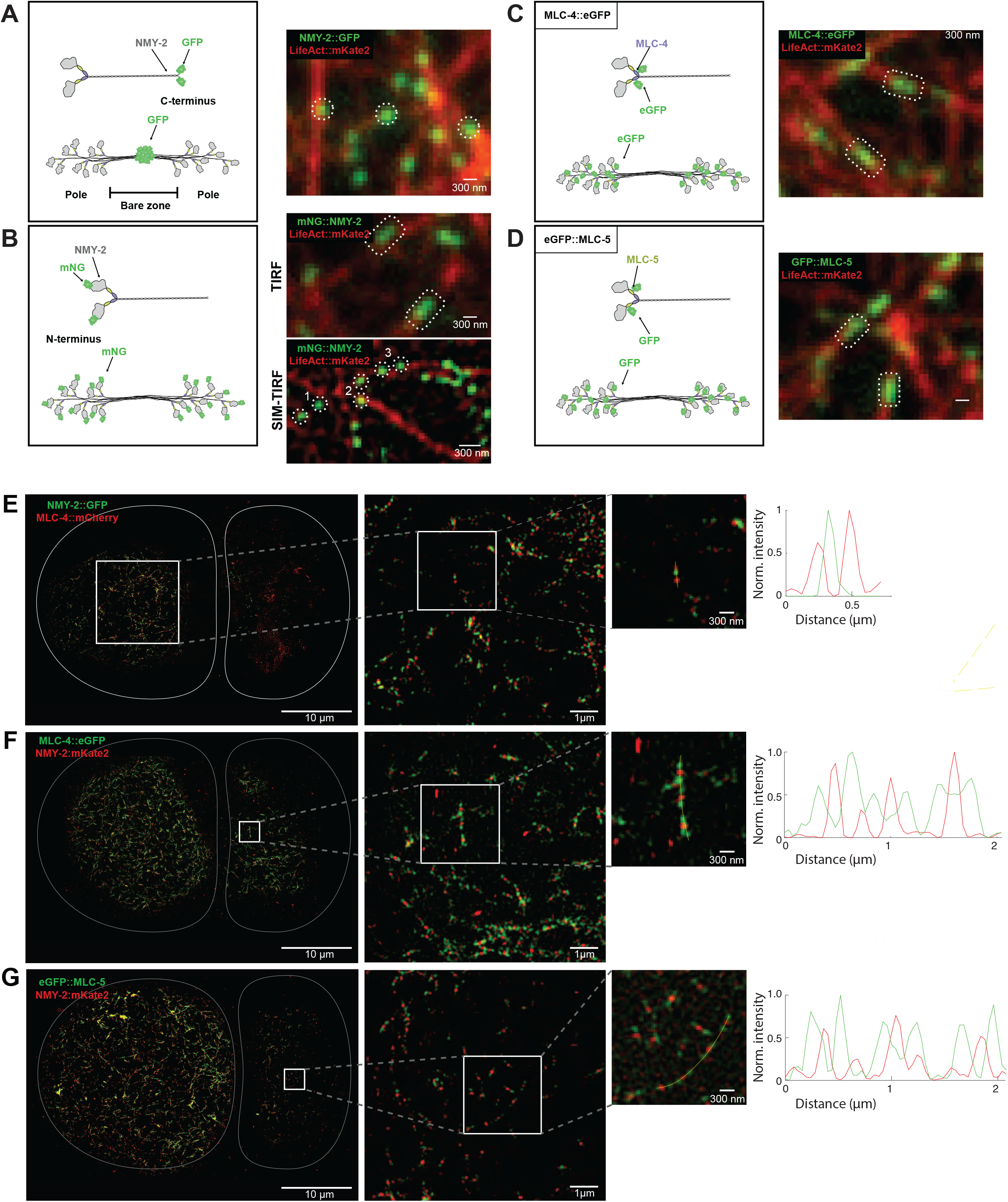
Strategic labeling of myosin subunits combined with TIRF imaging enables structure visualization of the myosin in early *C. elegans* embryo*s*. Left: Diagram illustrating subunits of the NMY-2 myosin monomer, composed of NMY-2 (myosin heavy chain), MLC-4 (regulatory light chain), and MLC-5 (essential light chain), strategically tagged at either the N- or C-terminus. Middle: Diagram illustrating the structure of myosin minifilament, functional contractile unit of force generation. Myosin minifilaments exhibit a bipolar architecture with a central bare zone separating the head clusters at each pole. The diagram indicates the relative position of fluorophores when the tagged subunits assemble into bipolar minifilaments. Right: Distinct projection patterns of myosin minifilaments after strategic labeling of myosin subunits and TIRF imaging. LifeAct::mKate2 (in red) was used as genetic background. **(A)** TIRF imaging of C-terminus fusion of myosin heavy chain with GFP, NMY-2::GFP, revealing a distinctive *dot-like* pattern at the cortex. Three representative minifilaments are highlighted, appearing as single spots in proximity to F-actin. **(B–D)** TIRF imaging of **(B)** N-terminus fusion of NMY-2 with mNeonGreen, mNG::NMY-2, **(C)** C-terminus fusion of MLC-4 with eGFP, MLC-4::eGFP, and **(D)** N-terminus fusion of MLC-5 with eGFP, eGFP::MLC-5 revealing a consistent *rod-like* pattern, in agreement with the bipolar architecture of myosin minifilaments. Representative minifilaments (n = 2 per panel) appear bipolarly associated with F-actin. **(E–G)** Two-color live SIM-TIRF imaging of embryos co-expressing two different fluorescently tagged myosin subunits during two-cell stage: **(E)** NMY-2::GFP and MLC-4::mCherry, **(F)** MLC-4::eGFP and NMY-2::mKate2, **(G) e**GFP::MLC-5 and NMY-2::mKate2. Individual bipolar minifilaments appeared as triplet dot patterns with either a (F, G) Green–Red– Green configuration or a (E) Red-Green-Red configuration, consistent with the relative structure of heavy and light chains within the bipolar assembly. Isolated single-color poles were not observed. Line-scan intensity profiles along the yellow lines drawn across individual minifilaments in higher magnification image, showing an average distance of approximately 150 nm between each pole and the minifilament center.

Conventional TIRF microscopy allows visualization of cortical NMY-2 assemblies as diffraction-limited puncta, but does not provide sufficient spatial resolution to reliably distinguish the two poles of individual bipolar minifilaments. In contrast, SIM-TIRF, enabling the separation and quantitative measurement of minifilament pole-to-pole distances that would otherwise appear as unresolved spots. Because cortical myosin structures are highly dynamic, the combination of high spatial resolution and temporal imaging stability provided by SIM-TIRF would be essential not only for resolving minifilament architecture but also for tracking their behavior during embryonic remodeling.

To better characterize the structural organization of myosin minifilaments, we turned to Structured Illumination Microscopy (SIM-TIRF) (Kner et al., 2009), which improves lateral resolution to approximately 100–120 nm. We observed that the structures we identified as “*rods*” using TIRFM improved drastically to reveal two clearly visible minifilament poles, producing a “*double dot*” pattern **(Fig. S1B**,**E-G, 1E-G)**. This increased spatial resolution enabled direct visualization of both mono-polar and bipolar interactions with the actin cortex, while also providing valuable information about the minifilament size (the distance from one pole to the other pole of the minifilament). Using the distance between the intensity peaks of each pole as a measurement proxy of the minifilament length, we found that the labeling of the three myosin components (mNG::NMY-2, MLC-4::eGFP and eGFP::MLC-5) exhibited a stereotypical size of approximately 300 nm, similar to the size found in other non-muscle myosin **(Fig. 1E-G)** (Niederman & Pollard, 1975; Billington et al., 2013; Beach et al., 2014; Melli et al., 2018).

To compare the localization between the different subunits, MLC-4 and MLC-5 relative to the heavy chain NMY-2, we generated strains combining one light chain with the heavy chain NMY-2 labeled at the tail domain, both chains pairs labeled with different colors (NMY-2::GFP; MLC-4::mCherry, MLC-4::eGFP; NMY-2::mKate2 and eGFP::MLC-5; NMY-2::mKate2). This strategy physically separated both labeled regions in space and confirmed their coassembly in bipolar minifilament. SIM-TIRF imaging revealed a clear, well-separated “*triplet*” minifilament structure **(Fig. 1E-G)**, demonstrating that MLC-4, MLC-5 and NMY-2 assemble to form stereotypical cortical minifilaments.

### 2. Characterizing the structure of myosin minifilaments in the early *C. elegans* zygote

As previous i*n vitro* studies showed that the number of motor domains within a minifilament can influence its mechanical properties, including force production and interaction with actin filaments, (Nagy et al., 2013; Melli et al., 2018), we next sought to quantitatively estimate the number of NMY-2 monomers within individual cortical minifilaments.

To estimate the number of motor domains per minifilament, we implemented a fluorescence intensity–based quantification strategy using TIRF microscopy. To this end, we compared the intensity of cortical myosin structures to that of a single-molecule reference, using CYK-1::eGFP as a calibration standard. To ensure that CYK-1::eGFP signals corresponded to single molecules, we performed partial photobleaching prior to measurement **(Fig. 2B, top panels)**. We then imaged NMY-2::eGFP under conditions minimizing photobleaching, where minifilaments appeared as diffraction-limited spots **(Fig. 2B, bottom panels)**, and corrected for local background. Under these assumptions, intensity calibrations yielded a median of ~30 NMY-2 motor domains, or 14-16 myosin hexamers per minifilament, or 7-8 hexamer per half filament **(Fig. 2C-D)**. To further characterize minifilament organization *in vivo*, we imaged cortical myosin during polarity maintenance at the anterior pole using SIM-TIRF microscopy, paired myosin minifilament extremities using a custom MATLAB pipeline, and measured distance between the paired centroids with subpixel precision **(Fig. 2A, S2A-D)** yielding a measurement precision of ~20 nm (HWHM, see also Methods and **Fig. S2E**,**F**). Using mNeonGreen::NMY-2 and MLC-4::eGFP at identical developmental stages, we measured similar overall distributions, with respective modes at 277 nm and 246 nm.

**Figure 2.**
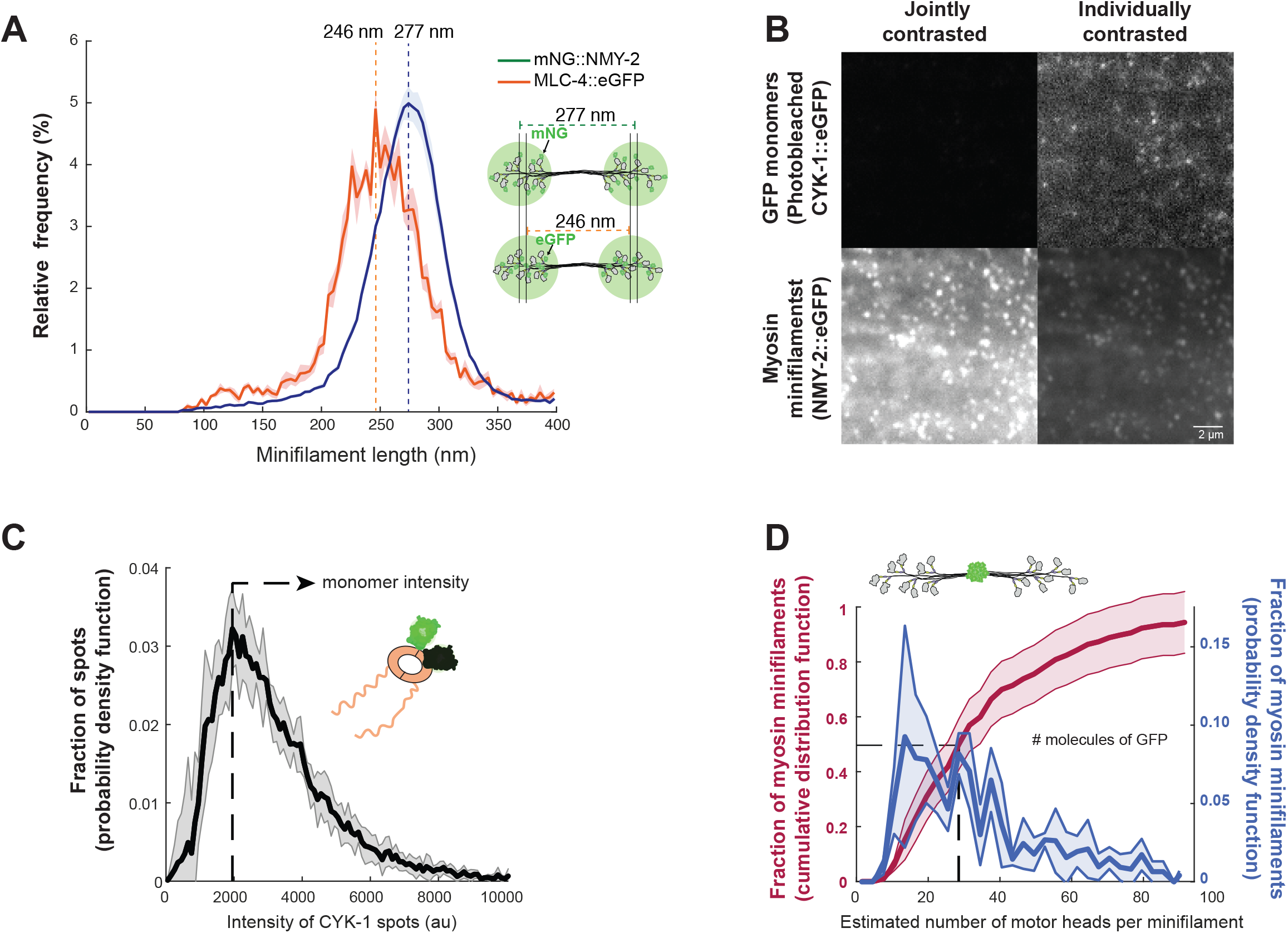
Characterizing the structure of myosin minifilaments in the early *C. elegans* zygote. **A)** Normalized relative frequency distributions of minifilament length measured using mNeonGreen::NMY-2 and MLC-4::eGFP strains under identical developmental stages. The mNeonGreen dataset shows a sharper distribution, consistent with improved photostability and tracking stability relative to eGFP. Dark solid lines represent the mean normalized distribution and shaded regions represent the standard error of the mean (SEM). Vertical dashed lines indicate modal peak positions. **(B)** TIRF microscopy images of NMY-2::eGFP (bottom) and CYK-1::eGFP (top), with image intensity scaling (left) and without (right). NMY-2::GFP was imaged for very briefly to avoid photobleaching, to preserve the fluorescence of myosin minifilament, while CYK-1::eGFP was heavily photobleached to ensure to that fluorescence intrinsic intensities corresponded to single eGFP. **(C)** Intrinsic intensities in camera counts for individual eGFPs (CYK-1::eGFP), N(embryos) = 5, n (measurements) = 30975. Vertical line corresponds to the mode of the distribution and was used as calibrating GFP intensity. **(D)** Estimated number of myosin motor domains per minifilament displayed as Cumulative Distribution Function (CDF, Red) or Probability Distribution Function (PDF, blue) N(embryos) = 5; n(measurements) = 607.

The superior photostability of mNeonGreen::NMY-2 in SIM-TIRF imaging led us to use as the reference dataset for ulterior quantitative analyses. Strikingly, in all measured conditions, the distributions displayed sharp modal peaks with left-skewed distribution with a steep drop-off after the mode, indicating that minifilament length is tightly constrained **(Fig. 2A)**, and displayed small spatiotemporal variations **(Fig. S2D** and data not shown).

These results demonstrate that the projected length of myosin minifilaments, despite minor developmental variations, exhibits a very narrow distribution, showing that the myosin minifilaments embedding in the cortex is predominantly parallel to the plasma membrane, potentially informative on cortical tension, cortex thickness, or minifilament penetration in the cortex.

### 3. NMY-2 minifilaments display previously hidden characteristic behaviors

As minifilaments could be observed as spatially isolated units, we decided to proceed to an analysis of their behavior, offering a window on how physiological regulation of myosin may alter minifilament cortical dynamics. We therefore used the high spatiotemporal resolution of SIM-TIRF microscopy to visualize both poles of individual minifilaments together with actin, and a customized semi-automated tracking pipeline to characterize their dynamics during polarity maintenance and the 2-cell stage, encompassing distinct cortical organizations and regulatory conditions.

Minifilaments were manually identified at their first appearance at the cortex and tracked until disappearance, while trajectory extraction and quantification were performed automatically, focusing on spatially isolated minifilaments to avoid trajectory mis-assignment (**see Materials and Methods**). Closer inspection revealed a consistent repertoire of five recurrent behaviors—static, walk, rotation, jump, and halo—observed across all developmental stages at a temporal resolution of Δt = 520 ms **(Fig. 3A–F, Videos)**. Similar behaviors were also observed at higher temporal resolution (Δt = 120 ms, **Videos**; see Supplementary Information), indicating that the classification is robust to imaging conditions. These behaviors could be broadly classified into actin-engaged behavior (*static, walk, rotation*), in which minifilaments remained fully or partially bound to F-actin, and detachment-associated behavior (*halo and jump*), reflecting transient unbinding from the cortical network. Interestingly these behaviors were reminiscent of the dynamics of myosin minifilaments observed in numerical simulations in works from our group as well as others.

**Figure 3.**
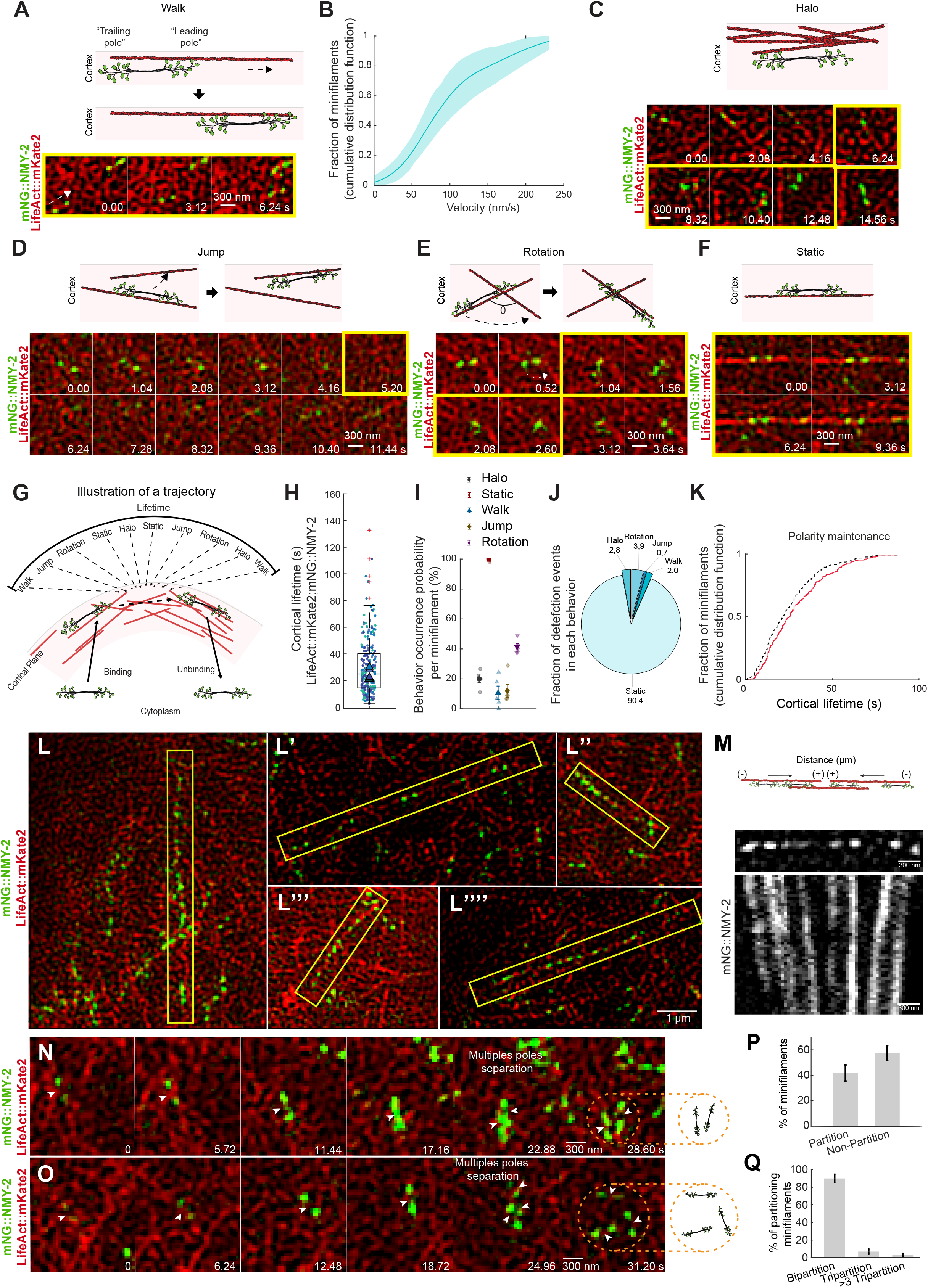
Super-resolution microscopy unveils myosin minifilament behaviors. **(A-L,N-O)** Characterization of myosin minifilament behaviors using SIM-TIRF microscopy of mNG::NMY-2; LifeAct::mKate2. **(A)** Top left: *walking* behavior illustration. Bottom left: Time lapse sequence showing a myosin minifilament moving from left to right. **(B)** Cumulative distribution of walking velocities. Quantitative analysis of the velocity for myosin minifilaments *walking* along F-actin was ~70-100 nm/s as measured during polarity maintenance (n = 28 minifilaments), with similar results during 2-cell interphase and 2-cell metaphase (not shown). Solid line shows the mean empirical cumulative distribution function (CDF) of walking velocities. Shaded area indicates the 95% confidence interval (CI) estimated by bootstrap resampling (1000 iterations) of all individual walking events, reflecting the uncertainty in the distribution. Walk events from embryos at the same developmental stage were pooled, as event numbers were very low or absent in some embryos. Some embryos contributed few or no walking events; the bootstrap CI captures the variability of the pooled events rather than variability between embryos. **(C)** Top: *halo* behavior illustration. Bottom: Time lapse sequence showing a myosin minifilament adopting a blurred appearance while moving rapidly yet remaining trackable. This behavior is interpreted as complete detachment from cortical F-actin followed by confined diffusion within the cortical actin meshwork or between the actin network and the plasma membrane. **(D)** Top: *jump* behavior illustration. Bottom: Time lapse sequence showing simultaneous detachment of both poles and rapid binding to a nearby F-actin filament. **(E)** Top: *rotation* behavior illustration. Bottom: Time lapse sequence showing detachment of one pole from F-actin followed by rebinding at a nearby actin filament, resulting in rotational reorientation. **(F)** Top: *Static* behavior illustration. Bottom: Time lapse sequence showing a myosin minifilament anchored to F-actin through both poles. This behavior may reflect alternating holding and leading motor activity within the bipolar assembly (schematic, top). **(G-K)** Characterization of the myosin minifilament lifetime at the cortex. **(G)** Diagram illustrating the cortical lifetime of a myosin minifilament, from cortical appearance (binding) to disappearance (detachment/unbinding). **(H)** Boxplots showing measured lifetimes (seconds) during polarity maintenance (IQR 14.4-40.2 s), N(embryos) = 5, n(minifilaments/embryo) = 45. **(I)** Probability that an individual minifilament exhibits at least one occurrence of a given behavior during its cortical lifetime. Percentages do not sum to 100 %, as individual minifilaments may exhibit multiple behaviors during a single lifetime. **(J)** Pie charts of the fraction of detection events per behavior, showing the distribution of minifilament behaviors across polarity maintenance. **(K)** Cumulative distribution function (CDF) plot quantifying halo-associated lifetime extension across the population. Dashed line: lifetime without halo; solid red line: lifetime with halo. Median lifetime extension was approximately ~5 s during polarity maintenance. N(embryos) = 5; n(minifilaments/embryo) = 45. Statistical analyses were performed as described in Methods. Significance levels: ns, P > 0.05; *P < 0.05; **P < 0.01; ***P < 0.001. **(L-M)** Accumulation of myosin minifilaments forming long, straight cables termed *proto-stress fibers* commonly observed during 2-cell interphase. **(L)** Myosin minifilaments in proto-stress fibers remained primarily static. **(M)** Proto-stress fibers often underwent global contraction, demonstrating that underlying actin bundles are at least partially antiparallel. **(N-Q)** Myosin minifilaments display partition-like behavior. Time-lapse sequence of a myosin minifilament undergoing partition in two **(N)** and three **(O)** “daughter” minifilaments. **(P)** Partition frequency in the myosin minifilament population (~42 % ± 6.25). N(embryos) = 5; n(bipolar minifilaments/embryo) > 45. **(Q)** Frequency minifilament bipartition (~90 % ± 4.7) and tripartition (~7 % ± 3) and more (3 % ± 1,9). N(embryos) = 5; n(minifilaments/embryo) > 12. Error bars represent 95 % confidence interval. Statistical analyses were performed as described in Methods. Significance levels: ns, P > 0.05; *P < 0.05; **P < 0.01; ***P < 0.001.

Strikingly, minifilament dynamics were dominated by the *static* behavior, which accounted for more than 87 % of the myosin minifilament detection events **(Fig. 3J)**. During this behavior, minifilaments remained largely immobile and anchored to F-actin through both poles **(Fig. 3F)** with episode durations ranging from ~18 to 23 s across developmental stages **(Fig. S4B)**. Consistently, nearly all minifilaments >98.7 % (671/675) exhibited at least one static episode during their lifetime **(Fig. 3I)**, establishing this state as the baseline mode of myosin interaction at the cortex. This state likely reflects the fact that both poles of the myosin minifilament are engaged in an interaction with actin filaments with opposing orientation, thus using ATP to generate force and set the network under tension.

In contrast, dynamic behaviors were comparatively rare and transient. During *walk*, minifilaments remained aligned with actin filaments while undergoing short, directional displacements. Interestingly, while NM2 minifilaments can move processively along actin cables *in vitro* (Melli et al., 2018), such processive motion is rarely reported *in vivo* outside specialized cells with highly bundled and parallel actin architectures, such as filopodia and sterocilia. In our system, the leading pole remained tightly associated with actin, whereas the trailing pole displayed increased positional fluctuations, suggesting asymmetric engagement during motility **(Fig. 3A)**. These events typically lasted ~5 s **(Fig. S4B)**, covered distances of ~300 nm **(Fig. S4C)**, and occurred at velocities ranging mostly from 70 to 100 nm/s **(Fig. 3B)**. *Walk* was observed in 11 to 16% of minifilaments across developmental stages (89/675 total; **Fig. S4F**), and accounted for less than 4 % of the detection events over all myosin minifilaments **(Fig. S4A)**.

*Rotation* behavior involved transient detachment of one pole followed by rebinding to the same or a different actin filament **(Fig. 3E)**. Approximately 38 to 51% of minifilaments exhibited at least one rotation event across developmental stages (292/675 total; **Fig. 3I, Fig. S4F**), although this behavior represented less than 5 % of the myosin minifilament detection events **(Fig. 3J)**. Rotation angles ranged between IQR 27.5-63 **(Fig. S4D)**, with exploration phases lasting ~1.5 s **(Fig. S4B)**.

*Jump* behavior, defined by the simultaneous unbinding of both poles followed by rapid rebinding to nearby actin filaments **(Fig. 3D)**, was rare, occurring in fewer than 12 to 15% of minifilaments across developmental stages (87/675 total; **Fig. S4F**) and corresponding to less than 1% of the total myosin minifilament detection events **(Fig. 3J)**. These events were brief (~1.5 s) **(Fig. S4B)** and involved diffusions of ~230 nm. Basic back-of-the-envelope calculations suggested that binding within in such narrow temporal and spatial proximity by another minifilament was extremely unlikely, driving us to attribute these rebinding events to the same minifilament, and the corresponding detachment to a local diffusion **(Fig. S4G)**.

Finally, minifilaments transiently lost a clear, well-defined bipolar appearance, transitioning to a blurred appearance, and displayed rapid motion, often before reappearing as discrete structures, a behavior we termed *halo* **(Fig. 3C)**, thus likely reflecting detachment events followed local diffusion of the minifilament within the laminated illumination optical sheet of HILO illumination plane. Halo events lasted ~3 to 4 s across developmental stages **(Fig. S4B)** and involved limited spatial diffusion ranging from ~260 to 370 nm across developmental stages **(Fig. S4C)**. While representing less than 5 % of the myosin minifilament detections events **(Fig. 3J)**, halo was relatively frequent at the population level, with 16 to 29% of minifilaments exhibiting at least one such event across developmental stages (147/675 total; **Fig. 3I, Fig. S4F)**.

Because the majority of minifilaments that went through halo reattached to the cell cortex, we wanted to see if halo could significantly increase the overall cortical lifetime of myosin minifilament **(Fig. 3H)**. To test if this impact was significant, we calculated the minifilament cortical lifetime including halo extension or without it. We found that halo increased the overall lifetime of minifilaments at the cortex during polarity maintenance on average by ~20%, from 21 s to 25 s **(Fig. 3K)**. These results also suggested that bipolar minifilaments are recycled as persistent, structurally intact assemblies.

During our analysis, we frequently observed groups of NMY-2 minifilaments aligned over large distances, with 3 to 8 minifilaments aligned with F-actin cables, spanning 2 to 5 µm, and with a median duration of ~14 s **(Fig. S3A-C)**, which we termed *proto-stress fibers* **(Fig. 3L,L’,L”,L’”,L”“)**. While myosin minifilaments participating in proto-stress fibers did not display clear preferential directional movement, proto-stress fibers often displayed clear contraction dynamics (**Fig. 3M)**, suggesting that these cable-like structures were under mechanical tension and that the underlying actin bundles displayed mixed polarity.

We also observed frequent partition-like events (Peterson et al., 2004; Shutova et al., 2014; Fenix et al., 2016; Beach et al., 2017; Najafabadi et al., 2022; Quintanilla et al., 2023) in which multiple NMY-2 minifilament signals emerged from the same cortical location occupied initially by a single detectable minifilament **(Fig. 3N,O)**. These events were defined based on the appearance of two or more spatially distinct minifilament structures originating from a pre-existing cortical site, without assuming the underlying molecular mechanism. Partition-like events were frequent, concerning (42 ± 6) % of analyzed minifilaments **(Fig. 3P)**, splitting minifilaments in two (90 ± 5) %, three (7 ± 3) % or more (3 ± 2) % **(Fig. 3Q)**.

Together, these observations indicate that cortical NMII organization is dynamically regulated across multiple spatial scales. Individual minifilaments can transiently engage in collective behaviors, including large-scale alignment into proto-stress fiber-like structures and local reorganization events that alter the number and spatial arrangement of detectable contractile units. Thus, cortical NMII organization emerges from dynamic transitions between individual minifilament states and collective configurations of force-generating units.

Together, these results define a consistent repertoire of minifilament behaviors across all developmental stages. Importantly, we observed only weak or not statistically significant differences between stages in the probability of experiencing each behavior (P > 0.05 for all comparisons) **(Fig. S4F,A-C)**, while myosin turnover itself changed **(Fig. S4E)**, supporting the conclusion that these dynamic states represent conserved properties of cortical myosin minifilaments during development.

### 4. Network stress is maximal when myosin exhibits static behavior

Our results thus show that minifilaments remain predominantly stationary, spending overall ~90% of their cortical lifetime in the static configuration (**Fig. 3J**), and bind to actin filaments for an average duration of ~20– 25 seconds (**Fig. 3H, Fig. S4E**, and **Fig. S4B** for static time). At the same time, the actin network is a dynamic material that continuously turns over, even during relatively stable phases such as the maintenance phase of the *C. elegans* first cell cycle, and remains under mechanical tension (Mayer et al., 2010).

To explore the interplay between myosin behavior and cortical tension generation, we used previously-established agent-based simulation (Jung, Murrell, and Kim, 2015, Costache et al, 2022). We used a three-dimensional box (5 µm x 5 µm x 100 nm) to simulated experimentally constrained regimes of F-actin filaments and myosin minifilaments kinetics (**Fig. 4A**). In particular we simulated nucleation, polymerizing, depolymerization and crosslinking of F-actin to mimic a cortex-like actin meshwork. Moreover, we introduce in the meshwork bipolar myosin minifilaments with 32 heads (matching our experimental results), each of which can switch from an unbinding state to a post power-stroke bound state and vice versa. By tuning the unbinding rate of single myosin head, we can affect the turnover of minifilaments.

**Figure 4.**
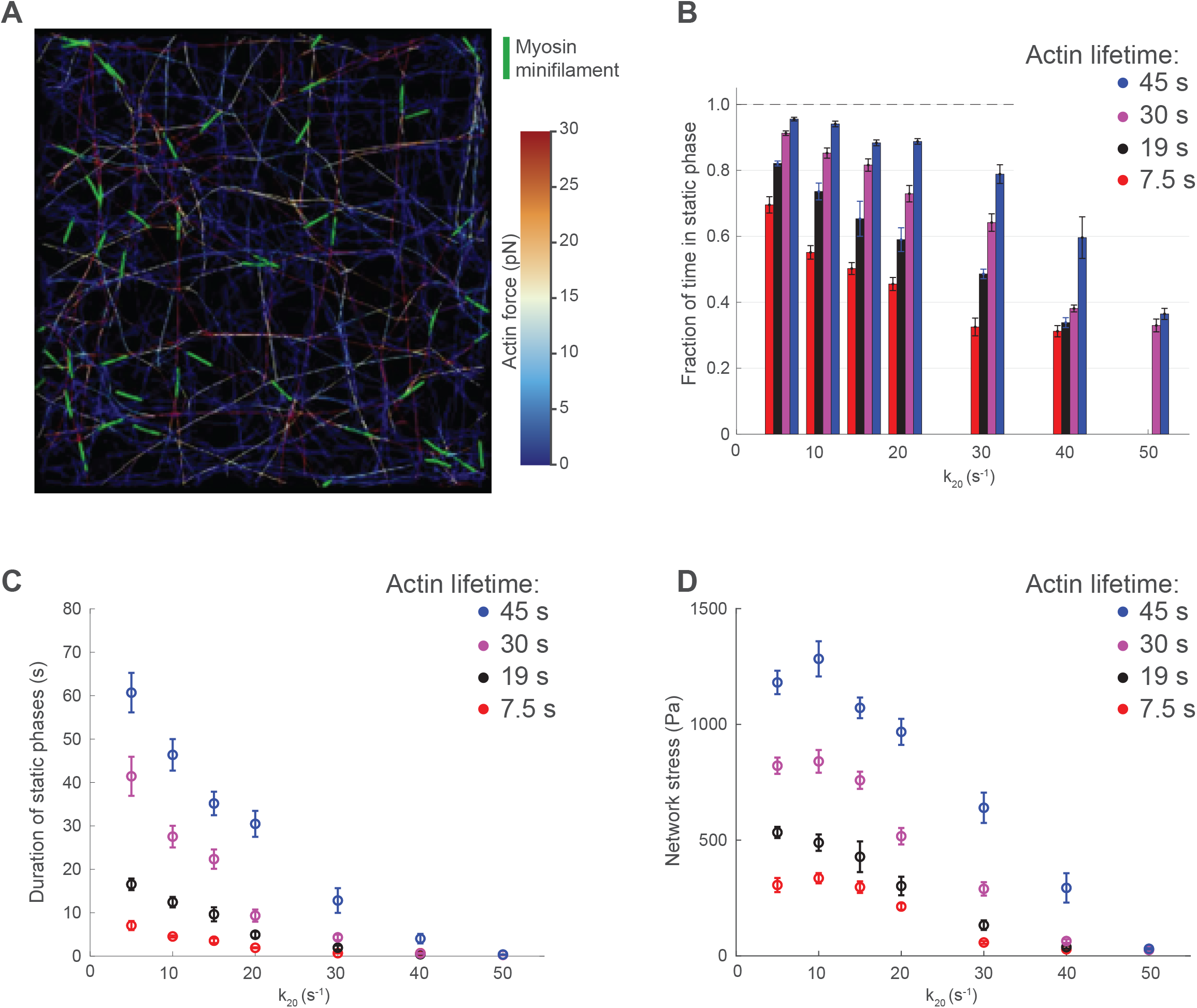
Network stress is maximal when myosin exhibits static behavior. **(A)** Frame extracted from of a simulated network of 5*µ*m x 5*µ*m x 100 nm containing 47 minifilaments with *k*_*20*_=10s^-1^ and average actin lifetime ~20s. Total duration of simulations is 150s. **(B)** Behavior of the fraction of time spent in static phases as function of the single head unbinding rate (*k*_*20*_) for 4 values of the average actin filaments lifetime. **(C)** Behavior of the static phases’ duration as function of *k*_*20*_ and for the same 4 values of the average actin filaments lifetime. **(D)** Behavior of actin network stress as function of *k*_*20*_ and for 4 values of the average actin filaments lifetime. Error bars are standard errors of the mean calculated on 6 independent simulation.

The cortical lifetime of the minifilament, as well as their kinetic behavior, is influenced by both cortical actin turnover and the intrinsic turnover of the minifilaments. Additionally, other parameters, such as the density of cross-linkers, myosin density, actin concentration, and actin filament length, may also affect myosins’ behavior and network stress. To simplify the analysis, we fixed the latter parameters and focused on the two primary dynamic cues of actin and myosin: (1) the unbinding rate of a single motor domain of the minifilaments from actin (called ***k***_***20***_) and (2) the cortical lifetime of actin filaments.

We focused on resulting duration of the static phases of the myosin (static time) and their fraction as read-out to compare with the experiment. As expected, the static time of minifilaments and its fraction decrease as ***k***_***20***_ increases (**Fig. 4B-C**). Additionally, actin turnover has a global effect, reducing the myosins’ static time. We then measured the behavior of network stress as a function of the single head unbinding rate and for different actin filaments turnover rate. Our results show that stress is a decreasing function of ***k***_***20***_ and exhibits a plateau at lower values of ***k***_***20***_. This plateau represents the maximal achievable stress at a given actin turnover rate and at a given myosin density. Furthermore, the plateau value itself is a decreasing function of actin turnover, which globally modulates the magnitude of stress in the network (**Fig. 4D**).

Given the static time duration of minifilaments and previous experimental data showing that the cortical lifetime of actin filaments is ~20 seconds (Robin et al. 2014), the stress regime of our biological system should be located in the plateau region, with ***k***_***20***_ **≤ 10 s**^**-1**^. In conclusion, since the minifilaments are highly static, the network achieves maximal stress.

### 5. NMY-2 motor activity controls myosin minifilament cortical lifetime at the cortex

Having established that cortical NMY-2 minifilaments exhibit a rich and structured repertoire of dynamic behaviors including transitions between stable binding, transient motility, and unbinding-associated states-we next asked how these dynamics are controlled at the molecular level. In particular, given that most behaviors reflect distinct modes of interaction with F-actin, we sought to determine whether myosin motor activity directly regulates minifilament cortical engagement and persistence.

The interaction between myosin and F-actin is governed by the ATP-dependent mechanochemical cycle of the motor domain, located within the S1 region. This domain, which includes both the motor and lever arm, plays a central role in actin binding, force generation, and stepping behavior. Mutations within this region are therefore expected to alter actomyosin interaction strength and kinetics, potentially impacting the dynamic behaviors described above (Kim et al., 2005; Uyeda et al., 2011).

To test this, we used myosin heavy chain mutants in the motor domain at the endogenous NMY-2 locus NMY-2^S250A^ and NMY-2^R718C^ **(Fig. 5A)** which were previously reported to alter cytokinesis duration by increasing the furrow initiation time and reducing the constriction rate (Osorio et al., 2019). To enable high-resolution analysis of minifilament behavior, these mutations were combined with MLC-4::GFP to report myosin localization and LifeAct::mKate2 to visualize F-actin, allowing direct comparison with the LifeAct::mKate2; MLC-4::GFP reference strain under SIM-TIRF imaging conditions.

**Figure 5.**
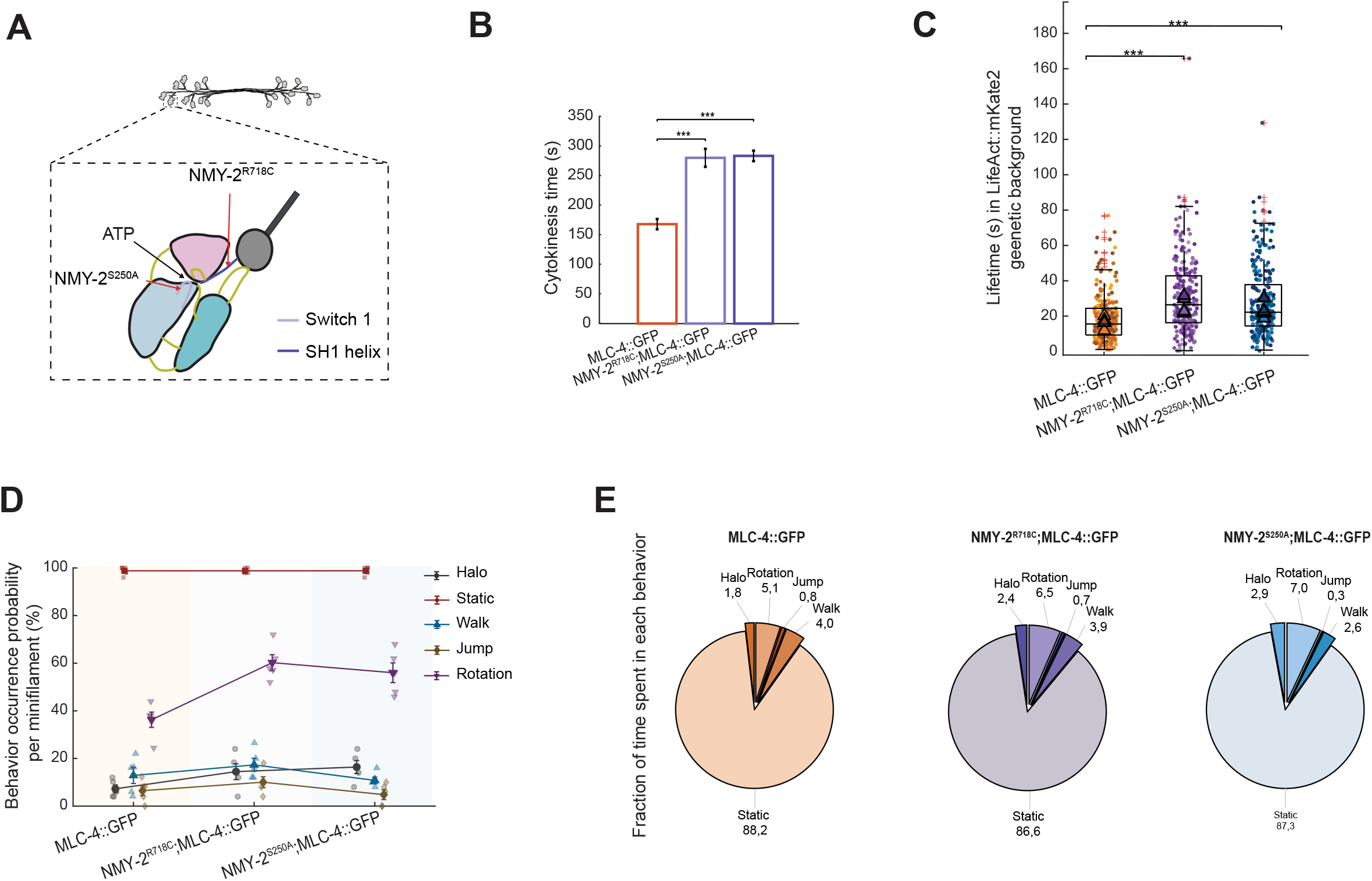
NMY-2 motor activity controls myosin minifilament cortical lifetime at the cortex. **(A)** Diagram illustrating mutation sites in the S1 motor domain of NMY-2. NMY-2^S250A^ is located in the Switch 1, while NMY-2^R718C^ is in the SH1 helix. **(B)** Bar plot showing cytokinesis duration in NMY-2^S250A^; MLC-4::GFP and NMY-2^R718C^; MLC-4::GFP mutants, compared to control MLC-4::GFP embryos (all strains in a LifeAct::mKate2 genetic background). Error bars represent ±95% confidence intervals (CI). Statistical analyses were performed as described in Methods. Significance levels: ns, P > 0.05; *P < 0.05; **P < 0.01; ***P < 0.001. **(C)** Boxplots displaying measured lifetime (seconds) for mutants NMY-2^S250A^; MLC-4::GFP ~20 s (IQR 14.6– 37.9 s; and NMY-2^R718C^; MLC-4::GFP ~23 s (IQR 16.4–42.9 s), and control MLC-4::GFP ~16 s (IQR 9.4–24.4 s). All strains co-express LifeAct::mKate2 for actin visualization). N(embryos) = 5 per genotype, n(minifilaments/embryo) > 45. Statistical analyses were performed as described in Methods. Significance levels: ns, P > 0.05; *P < 0.05; **P < 0.01; ***P < 0.001. **(D)** Probability that an individual minifilament exhibits at least one occurrence of a given behavior during its cortical lifetime in control MLC-4::GFP, NMY-2^R718C^; MLC-4::GFP, and NMY-2^S250A^; MLC-4::GFP embryos. Multiple behaviors may occur within a single lifetime; therefore, percentages do not sum to 100 %. **(E)** Pie charts of time fractions per behavior (number of detection events in the behavior over total number of detection events), showing the distribution of minifilament behaviors during 2-cell interphase in control MLC-4::GFP, NMY-2^R718C^; MLC-4::GFP, and NMY-2S^250A^; MLC-4::GFP embryos. Percentages represent the proportion of total behavioral events per genotype; values within each pie chart sum to 100 %.

Since the addition of a fluorescent label near the S1 region could potentially interfere with myosin function, we first validated that these strains recapitulated previously reported phenotypes by Osorio et al., 2019. Using brightfield microscopy, we measured the duration of the first cytokinesis and observed a significant increase in both mutant backgrounds (NMY-2^S250A^; MLC-4::GFP; LifeAct::mKate2 and NMY-2^R718C^; MLC-4::GFP; LifeAct::mKate2) compared to the corresponding control strain (MLC-4::GFP; LifeAct::mKate2), consistent with prior reports (Osorio et al., 2019) **(Fig. 5B)**, confirming previously documented functional impact of the mutations.

We then quantified cortical minifilament dynamics in these mutant backgrounds. Strikingly, both mutations led to a significant increase in minifilament cortical lifetime. In NMY-2^R718C^, lifetime increased by ~7 s relative to control, from ~16 s to ~23 s (***P < 0.001). Similarly, in NMY-2^S250A^ lifetime increased by ~4 s, from ~16 s to ~20 s (***P < 0.001; **Fig. 5C**).

In addition to these changes in lifetime, we observed alterations in specific dynamic behaviors. While most behaviors remained largely unchanged across conditions, motor perturbations produced selective effects on transient cortical states. Halo behavior showed a modest but statistically significant modulation across conditions, indicating altered frequency of detachment-associated events **(Fig. 5D)**.

In contrast, rotation behavior was more strongly affected at the single-event level. Although the fraction of time spent in rotation increased only modestly in both mutants (from ~5.1 % in control to ~6.5 % in NMY-2^R718C^ and ~7.0 % in NMY-2^S250A^; **Fig. 5E**), the probability that a minifilament underwent at least one rotation event during its lifetime increased substantially, from ~33 % in control conditions to ~57 % in NMY-2^R718C^ and ~60 % in NMY-2S^250A^ **(Fig. 5D)**, suggesting that the occurrence of this behavior might simply scale with the cortical residence time.

### 6. Myosin photoconversion reveals marginal minifilament cytoplasmic dissolution and key role for recycling of myosin minifilament following unbinding from the cortex

Our previous results showed that myosin minifilaments persist following unbinding during *halo* and *jump* behavior, with little apparent disassembly. These observations led us to hypothesize that minifilaments undergo only limited disassembly upon unbinding, instead retaining their organization over extended periods of time. To test this hypothesis, we turned to a pulse-chase strategy using photoconversion using a Dendra2-Myosin.

We first generated a CRISPR knock-in strain expressing Dendra2-Myosin at the endogenous genomic locus (Gurskaya et al., 2006). Consistent with previous reports in *C. elegans* (Graner et al., 2008; Griffin et al., 2011; Rosu & Cohen-Fix, 2017), we validated the efficiency and spatial specificity of photoconversion under our experimental conditions. To this end, we first performed full-field cortical photoconversion prior to pulse-chase experiments. Upon 405 nm illumination, green cortical minifilaments were rapidly converted into red fluorescent structures, with near-complete loss of green signal and concomitant appearance of red fluorescence. Within ~30 s, new green minifilaments repopulated the cortex, while photoconverted red minifilaments redistributed, indicating exchange with a cytoplasmic pool (**Fig. 6A, Video see Material and methods**). These observations validated the conditions used for subsequent pulse-chase experiments.

**Figure 6.**
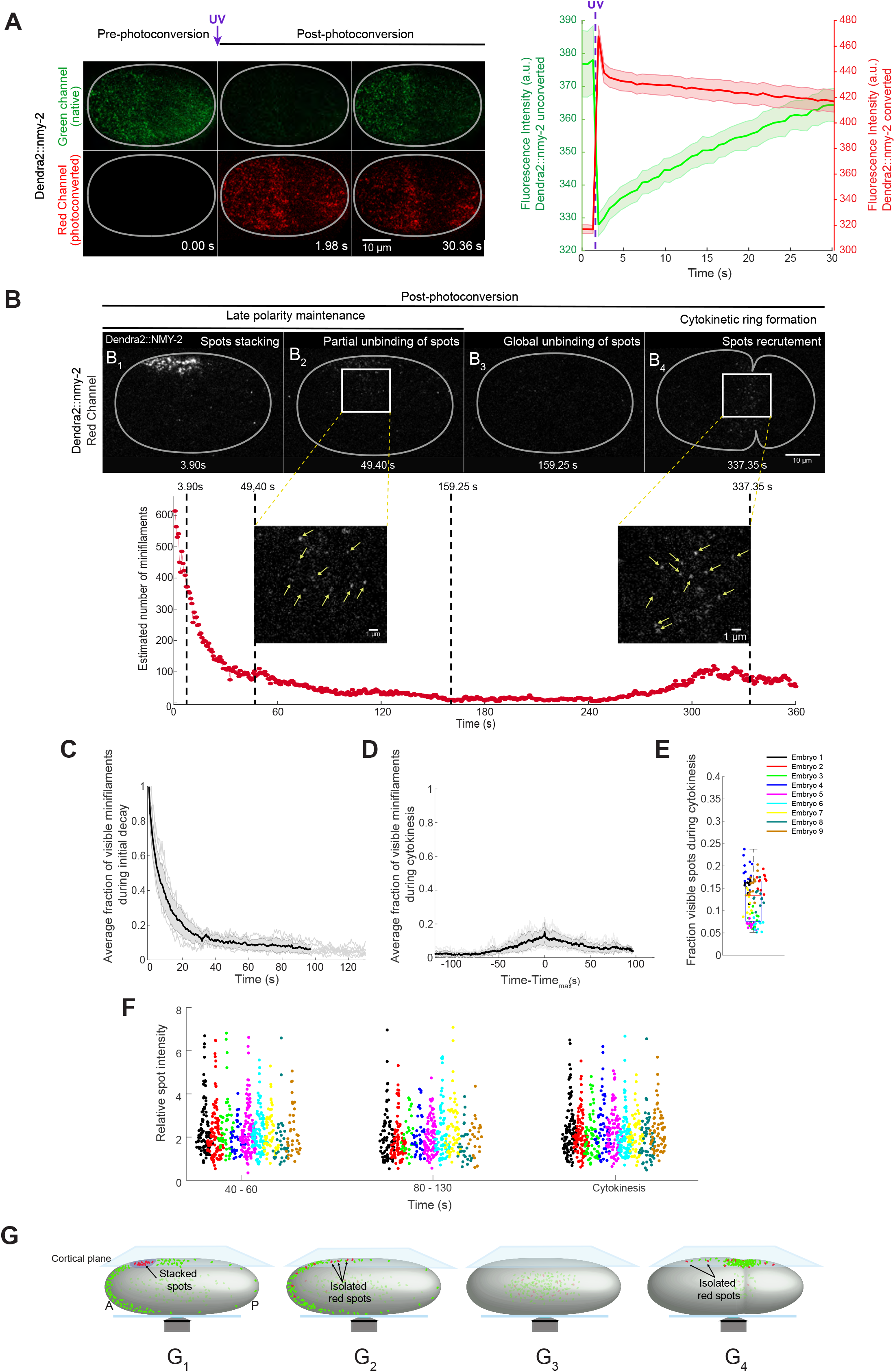
Photoconversion pulse-chase analysis reveals that myosin minifilaments persist and are not disassembled upon cortical detachment. **(A)** Validation of Dendra2 photoconversion efficiency. Prior to UV exposure, red fluorescence was negligible, while strong green cortical signal corresponding to Dendra2::NMY-2 minifilaments was observed. Immediately after photoconversion (t ≈ 2 s), green fluorescence decreased sharply, accompanied by a concomitant increase in red signal, indicating efficient photoconversion of cortical minifilaments. At later time points (t ≈ 30 s), green fluorescence progressively recovered at the cortex, while red signal gradually decreased, consistent with turnover and exchange of photoconverted minifilaments with a cytoplasmic pool. Left: Time-lapse sequence of a representative full-field cortical photoconversion experiment of Dendra2::NMY-2. Right: Quantification of fluorescence intensity (a.u.) of non-photoconverted (green) and photoconverted (red) Dendra2::NMY-2 over time. Thick lines represent averages, shadows represent SEMs. N(embryos) = 9. **(B)** Top: Time lapse sequence of a representative Dendra2::NMY-2 pulse-chase experiment. Analysis performed from polarity maintenance (pronuclear meeting) until furrow formation. **(B1)** Representative frame of an embryo 3.90 s after photoconversion during the polarity maintenance. **(B2-B4)** Images of the photoactivated signal at 49.40 s **(B2)**, 159.25 s **(B3)**, and 337.35 s after photoconversion **(B4)**. The initial photoactivated patches progressively separate into smaller spots **(B2)**. Spots disappear from the cortex to the cytoplasm during clearing phase **(B3)** and reappear during the furrow formation. **(B4)** Bottom: Estimated number of minifilaments corresponding to the sequence shown in **(B1–B4)**. Minifilaments number progressively decreases after photoconversion reaching a minimum during clearing phase. Subsequently the number of minifilaments increases, peaking at furrow formation before cell division. Vertical lines indicate the frames displayed in panels **(B1–B4)**. At 49.40 s and 337.35 s, magnified views of the regions indicated by rectangles in **(B2)** and **(B4)**. Results for embryo collection are presented in **Fig. S5B. (C)** Average decay of the fraction of visible spots (minifilaments) as a function of time from photoconversion (pronuclear meeting of maintenance phase) to the end of maintenance phase. N(embryos) = 9. The fraction progressively decreases after photoconversion, reaching a pseudo-plateau that decease, corresponding to the polarity maintenance. **(D)** Average fraction of visible spots measured around the signal peak at furrow formation, all the curves have been aligned respect to the frame where the fraction is maximal (time_max_). **(C-D)** Thick black line represents the average, the gray shadow represents the standard deviation and light gray thin curves the individual embryos behaviors. **(E)** Fraction of visible spots estimated at the furrow (11 frames around the maximum during furrow formation). See also **Fig. S5C**. N(embryos) = 9. The fraction ranges from 5 % to 20 %, with a median value of ~13 %. **(F)** Distributions of the relative total fluorescence intensity of spots. N(embryos) = 9. Intensities were measured at two early time points (45 s and 60 s), at two time points after the first minute (~80 s and ~130 s), and at two time points near furrow formation. The relative intensity is defined as *I*/(2*< I*_*eq*_*>* _*time>60s*_), with *I* the total intensity and *< I*_*eq*_*>* _*time>60s*_ is the time average of the intensity starting from the first minute till the end of each movie. Statistical analyses were performed as described in Methods. Significance levels: ns, P > 0.05; *P < 0.05; **P < 0.01; ***P < 0.001. **(G)** Schematic model illustrating the dynamics of photoconverted and non-photoconverted minifilament spots between the cortex and cytoplasm from late polarity maintenance to furrow formation. **(G1)** Immediately after UV photoconversion within the ROI, individual spots cannot be resolved due to local stacking of photoconverted minifilaments. **(G2)** Within seconds, discrete red spots become detectable as minifilaments disperse into the cytoplasm and redistribute along the cortex. **(G3)** During the clearing phase, both red (photoconverted) and green (non-photoconverted) cortical spots undergo global redistribution into the cytoplasm. This stage is critical for our analysis, as previous studies have proposed that cytoplasmic monomer exchange or de novo minifilament assembly occurs during this phase. If disassembly into monomers were occurring, red spots generated prior to clearing would be expected to markedly diminish or disappear and would not persist until furrow formation. **(G4)** In contrast, we observe that the integrated intensity of photoconverted minifilaments measured 49,4 s after activation is comparable to the intensity of red spots detected during furrow formation, supporting the persistence of pre-existing minifilament structures. N(embryos) = 9.

During the pulse-chase experiment we photoactivated Dendra2-Myosin in a small circular region of the cortex using a 405 nm laser in TIRF mode **(Fig. 6B, Video, see Material and methods)**. As expected, we could activate Dendra2-Myosin as a discrete cortical region with little/no ectopic activation, allowing us to track the dynamics of photoconverted myosin over time. Initially, photoconverted myosin appeared as confluent aggregates, or patches **(Fig. 6B**_**1**_), which decreased in size within the first ~40-60 s **(Fig. S5A)** following photoconversion, then reaching a plateau lasting for several minutes **(Fig. S5A)**. The average size of the detected spots thus reached a stable value that was maintained throughout the entire movie, irrespective of the number of spots present and total cortical intensity **(Fig. S5A)**. We estimated the lifetime of individual myosin minifilaments at the cortex to be around ~20-25 s and, as a consequence of minifilament progressive unbinding from the cortex, patches progressively diluted until eventually only individual minifilaments remained visible.

We assume a that the average constant value of spots after the initial decay correspond to the average constant minifilament size throughout the experiment (insets **Fig. 6B_1_-B_4_)**. Indeed, spots size appear similar at the beginning and at the end of the movies (insets **Fig. 6B_2_-B_4_)**. We then then rescaled divided our intensity as by the average minifilament intensity obtaining the number of minifilaments present at the cell surface **(Fig. 6B, red symbols)** and then plotted as a fraction of minifilament present initially during the first 80 s following photoconversion **(Fig. 6C)**. While between 150 s and 250 s following photoconversion, we observed no cortical spots **(Fig. 6B**_**3**_), furrow formation was associated with the robust reappearance of multiple spots **(Fig. 6B**_**4**_, **Fig. 6D, Fig. S5B, C)**, ranging from 5 to 25 % of the initially photoconverted minifilaments **(Fig. 6D-E)**, with a similar intensity as observed previously **(Fig. 6F)**. Taken together, these observations show that a fraction of the initially photoactivated minifilaments persists as stable units until the first cell division.

Following this observation, we wanted to quantitatively infer from these results the levels of dissolution fraction throughout our observation window. To this end, we designed a simple kinetic model where the cortical minifilaments exchanged with a cytosolic reservoir (see *Materials and Methods* for model details). From this simple model, we expected to observe about 10 % of the initial photoactivated minifilament pool recruited at the furrow in the visible view during cytokinesis. Strikingly, our measurements showed fractions ranging from (5 to 20) %, with an average of (15 ± 6) % **(Fig. 6D, E)**, a result fully consistent with a *no dissolution* model.

In conclusion, we showed that a significant fraction of the initially estimated minifilaments reappear at the furrow during cytokinesis, consistent with the persistence and reuse of the minifilaments, and supporting that dissolution of minifilaments into monomers upon unbinding from the cortex remains marginal over periods of >10 min in the early embryo **(Fig. 6G)**.

## DISCUSSION

Non-muscle myosin II (NMII) is a central generator of cortical tension during embryogenesis, yet the organizational principles governing individual cortical minifilaments *in vivo* remain poorly understood. Previous studies have largely emphasized highly dynamic cortical remodeling associated with pulsatile RhoA signaling, transient cortical recruitment of myosin, and rapid changes in cortical myosin accumulation during embryogenesis (Munro et al., 2004; Robin et al., 2014; Prigent Garcia et al., 2026). However, whether these fluctuations reflect continuous de novo assembly and disassembly of individual minifilaments or instead changes in the cortical engagement and redistribution of persistent preassembled structures has remained unclear. Understanding how the early embryo cytoskeleton maintains mechanical activity while remaining highly dynamic requires resolving not only the organization of myosin assemblies, but also their behavior and lifetime within the cortex. Combining super resolution imaging with single-particle tracking, we showed that embryonic cortical NMII is organized as persistent bipolar minifilaments. Here we discuss the measured features of minifilaments and how the different modes of cortical engagement may contribute to force generation in the developing cortex.

The dimensions of cortical minifilaments measured here were broadly consistent with previous ultrastructural and super-resolution studies of canonical NMII bipolar assemblies, which exhibit relatively stereotyped nanoscale organization and molecular packing (Niederman & Pollard, 1975; Billington et al., 2013; Beach et al., 2014; Fenix et al., 2016). Our analyses extend these observations to the early embryonic cortex in vivo and show that minifilament dimensions **(Fig. S2D)** and inferred monomer numbers (data not shown) remain relatively constrained across developmental stages. This stability argues against extensive fragmentation or stochastic de novo assembly and instead supports the existence of persistent contractile units that retain their bipolar organization while dynamically changing their cortical interactions. Although we detected small variations in projected minifilament length, these may reflect fine-scale changes in filament positioning or orientation within the cortex, potentially influenced by cortical architecture or myosin regulatory pathways (Truong Quang et al., 2021; Amano et al., 1996; Matsumura, 2005; Vicente-Manzanares et al., 2009). Thus, while embryonic minifilaments maintain a remarkably stable overall architecture, their mechanical configuration may remain locally tunable.

A striking feature of our observations was the formation of aligned and stacked minifilament structures within the embryonic cortex. Similar stacking has been described in stress fibers and contractile arrays, where neighboring bipolar filaments form larger force-generating assemblies (Fenix et al., 2016; Hu et al., 2017). Proposed mechanisms include actomyosin flows and filament collisions, templating by actin architecture, and self-organization driven by contractile forces and nematic ordering (Fenix et al., 2016; Hu et al., 2017; Burnette et al., 2014; Murrell & Gardel, 2012; Koenderink & Paluch, 2018). Our observations are compatible with these mechanisms and, importantly, show that higher-order minifilament alignment can emerge within the highly dynamic embryonic cortex, despite the absence of mature stress fibers. These transient stress-fiber-like structures may locally reinforce cortical tension, promote directional force propagation, or stabilize mechanically polarized domains during rapid developmental remodeling (Tojkander et al., 2011; Burridge & Wittchen, 2013). Together, these observations highlight the mechanical heterogeneity of the embryonic cortex, which can organize contractile forces across scales from individual minifilaments to larger supramolecular assemblies.

Our dynamic analyses further revealed that cortical minifilaments transition between multiple behavioral states, including static anchoring, directional walking, rotational exploration, transient diffusion, and rapid rebinding events. Strikingly, the static state accounted for the overwhelming majority of cortical lifetime, indicating that most minifilaments spend prolonged periods mechanically engaged with the cortical actin network. As a result of the predominant static behavior of minifilaments, the actomyosin cortex maintains a state of sustained mechanical stress. Our simulations further demonstrate that, considering the turnover rates of actin and myosin, the network operates near its maximum possible stress, constrained by motor density and actin filament connectivity. In essence, despite the dynamic nature of cytoskeletal turnover, this prolonged motor engagement is crucial for sustaining contractile force within the cortex.

The motor-domain mutants analyzed here provide mechanistic insight into how ATP-dependent motor activity regulates cortical minifilament dynamics. The S1 region of myosin plays a central role in actin binding, force production, and mechanochemical cycling, and perturbations within this domain are expected to alter actomyosin interaction kinetics (Kim et al., 2005; Uyeda et al., 2011). Consistent with this expectation, both NMY-2S250A and NMY-2R718C increased cortical lifetime, suggesting that altered motor kinetics prolong cortical association rather than destabilizing it. One possibility is that these mutations modify effective dissociation rates from actin or alter the duty ratio of the motor, thereby extending the persistence of force-bearing interactions at the cortex. Rotation behavior was particularly sensitive to perturbations of the motor domain. Because these mutations affect regions involved in ATP-dependent actin binding and force generation, the increased probability of rotational events likely reflects altered coordination between the two poles of the bipolar minifilament. Rather than functioning as perfectly synchronized force-generating units, mutant minifilaments may experience transient imbalances in actomyosin engagement, where one side of the minifilament remains stably attached while the opposite side undergoes delayed detachment or inefficient rebinding. Such asymmetries would increase the probability of local pivoting around a single anchoring point and promote repeated exploration of neighboring actin filaments. Importantly, this interpretation suggests that rotation is not merely a passive geometric behavior but may represent an active search mechanism through which minifilaments reorganize their local mechanical connectivity within the cortical network. The simultaneous increase in cortical lifetime further suggests that motor perturbation does not simply weaken cortical association. Instead, could alter transitions through the mechanochemical cycle, increasing the persistence of individual actomyosin interactions and thereby stabilizing cortical residence while favoring repeated local rearrangements. Together, these observations indicate that motor activity regulates not only the magnitude of force generation but also the dynamic balance between stable engagement and exploratory remodeling within the embryonic cortex.

Our findings also have implications for understanding embryonic cortices as active materials. Current active matter descriptions often model actomyosin networks as collections of highly dynamic motors interacting stochastically with actin filaments. However, the persistence and reutilization of cortical minifilaments observed here suggest a distinct physical regime in which long-lived force dipoles repeatedly engage and disengage from a dynamically evolving actin meshwork. Importantly, maintaining a reusable pool of preassembled minifilaments may confer major advantages during early embryogenesis, where cortical mechanics must remain simultaneously robust, adaptive, and rapidly responsive over developmental timescales of seconds to minutes.

Several limitations should nevertheless be considered. Although our imaging approach achieves high spatial and temporal resolution, it does not directly resolve individual motor-head interactions or subunit exchange occurring below the detection threshold. Likewise, while our observations strongly support reutilization of intact minifilaments, they cannot formally exclude limited molecular turnover within persistent bipolar assemblies. Finally, the mechanistic models proposed here remain inferential and will require additional experiments combining live imaging with simultaneous readouts of cortical signaling and actomyosin mechanics.

In summary, our work identifies the embryonic cortical actomyosin network as a highly dynamic yet structurally persistent mechanical system organized across multiple spatial scales. Individual myosin minifilaments behave as persistent bipolar assemblies capable of self-organizing into aligned and stacked higher-order structures while dynamically transitioning between multiple cortical interaction states. Together, these findings support a model in which embryonic cortical mechanics emerge not from continuous reconstruction of actomyosin structures, but from regulated cycles of engagement, redistribution, and reutilization of persistent force-generating units.

## MATERIAL AND METHODS

### Experimental model

All *C. elegans* strains used in this study were cultured and maintained using standard methods described previously (Brenner, 1974). All strains were fed with *E. coli* OP50, grown on NGM agar plates in the dark at 20°C inside incubators unless otherwise stated. Strains used in this study are listed in **Table S1**. Transgenic strains were constructed by CRISPR/Cas9 genome editing techniques, following previously published protocols (Dickinson et al., 2013; Dickinson & Goldstein, 2016). Briefly, the cocktail of DNA constructs was injected into N2 adult *C. elegans* germline syncytium. The cocktail of plasmids comprised a repair template, pDD162 (Dickinson et al., 2013), a SG coding plasmid and a suitable co-injection marker encoding plasmid such as pRF4 or pCFJ90 (Frøkjær-Jensen et al., 2008).

The homologous repair template used in mlc-4 and nmy-2 fluorophore knock-in transgenesis were made by the stitching of purified gene-specific PCR fragments on desired plasmid backbone (vector from Thermo Fisher Scientific or pJJR83 from Addgene) using the Gibson Assembly Cloning kit (New England Biolabs, Inc.). The homologous repair template usually contains >1500 bp and >700 bp genomic DNA flanking to the site of fluorophore insertion into the gene. The DNA sequence of fluorophores mCherry and mNeon were taken from pJJR83, and dg397 plasmids respectively. For the construction of a gene-specific SG plasmid, pML2840 (Suman et al, 2019) was used as a backbone and gene specific 20 bp CRISPR RNA equivalent DNA sequence was incorporated via MEGAWHOP extension PCR method (Lee et al., 2010). The details of primers, SG sequence and plasmids are indicated in **Table S2, S3 and S4** respectively. All following steps were performed as described in Dickinson et al, 2013 & Dickinson et al, 2016. Transgenic lines were outcrossed with N2 strain and generally maintained at 20°C. All the transgenics line were further verified by PCR based genotyping and DNA sequencing.

### Total Internal Reflection Fluorescence Microscopy (TIRFM)

TIRF imaging was performed as previously described (Robin et al., 2014). Young adult hermaphrodite worms were dissected in 3 μl of water supplemented with ~100 polystyrene beads (15.6 ± 0.03 μm diameter; Bangs Laboratories, #NT29N) on 3-well glass slides and mounted under 22 × 22 mm high-precision coverslips. All imaging experiments were performed at 21°C.

Embryos were illuminated using solid-state lasers (maximum capacity 300 mW), with laser output set to 30% of the maximum (corresponding to 90 mW). Laser intensity was controlled via the microscope software, where 100% corresponds to the full transmitted power under these conditions. Laser power was measured at the objective under TIRF illumination conditions, with the incidence angle set using NIS-Elements (angle = 0) and corresponded to approximately 12.4 mW (561 nm), 4.6 mW (488 nm), and 2.4 mW (405 nm).

Fluorescence emission was collected using an Apo TIRF 100× oil-immersion DIC N2 objective (Nikon) with a numerical aperture (NA) of 1.4, coupled to an *Andor iXon 897* EMCCD camera with a 1.5× Optovar and Nikon Perfect Focus System, resulting in a final pixel size of 107 nm. Image acquisition parameters (angle, laser intensity, and exposure) were controlled using NIS-Elements software (version 4.50, Nikon). For all experiments, the laser incidence angle was empirically adjusted to maximize signal intensity while maintaining uniform illumination across the field of view. For two-color datasets, image acquisition of each channel was performed sequentially.

For Fig. 1, imaging of mNeonGreen and mKate2 was performed using 488 nm and 561 nm excitation, respectively, with 50 ms exposure time at 15 % laser power, and an acquisition rate of 10-20 frames per second. For Fig. 6A, unconverted Dendra2 was imaged using 488 nm excitation (50 ms exposure, 11 % laser power), whereas photoconverted Dendra2 was imaged using 561 nm excitation (50 ms exposure, 11 % laser power), with an acquisition interval of 660 ms. For Fig. 6B, imaging was performed using 561 nm excitation (50 ms exposure, 30 % laser power) with a 130 ms acquisition interval. To minimize unintended photoconversion, the green form of Dendra2::NMY-2 was visualized using a 488 nm laser at low power (8 %) and an acquisition interval of 130 ms.

### TIRF-SIM microscopy

Embryos were prepared in slides following the same protocol as for TIRF microscopy. Images were collected on a DeltaVision OMX SR imaging system (OMX version 4.3.9379-1) from GE Healthcare (Velizy, France) with an Olympus PlanApo N 60× 1.42 NA oil immersion objective, using the immersion oil (Cargille laboratories Inc), providing an *n*D (at 21°C and 589.3 nm) = 1.5140 ± 0.0002 to minimize spherical aberrations. For mNeonGreen and mKate2 fluorescent probe, imaging was acquired using excitation at 488 nm during 10 ms (15 %) and 568 nm during 10 ms (20 %), respectively.

For all samples, SIM images of 512 × 512 pixels were acquired at single z-plane with perfect focus mode. Each TIRF-SIM image is a result of the average reconstruction of 9 raw images (three phases at three angles) and its final pixel size was 40 nm. The image quality was evaluated by the SIMcheck (Ball et al., 2015), a suite of Fiji plugins (Schindelin et al., 2012). Finally, images were aligned correcting chromatic aberration by the SoftWoRx Explorer Software (Applied Precision Inc) which enables a resolution enhancement in the order of 100 nm.

Based on its superior photostability, signal continuity, and reduced photobleaching under SIM-TIRF imaging conditions, mNeonGreen was used as the primary reference dataset for all quantitative analyses requiring sub-diffraction resolution of pole-to-pole distances. In contrast, eGFP was used for lower spatial precision.

### Determination of the NMY-2 minifilament length

Single-minifilament tracking was performed using a MATLAB implementation of the Crocker–Grier particle tracking algorithm (Crocker & Grier, 1996) provided by the Kilfoil lab. Fluorescent spots corresponding to the poles of myosin minifilaments were detected with subpixel precision and subsequently linked across frames. To identify genuine doublets, we implemented a custom MATLAB pipeline based on mutual nearest-neighbor pairing within each frame. Candidate pairs were required to satisfy two criteria: (i) a nearest-neighbor distance within a defined physical range, and (ii) a second-nearest-neighbor distance above a threshold to exclude ambiguous or crowded detections.

For the tracking of myosin in FBR159 *gesIs001[Pmex-5::LifeAct::mKate2::nmy-2(3’UTR); unc-119(+)]; nmy-2(cp141[mNG^3xFlag::nmy-2])I* strain, particle detection was performed using *mpretrack* with parameters: masscut = 450, minimum intensity = 30, and featsize = 3. Doublet identification used nearest-neighbor distances of 2– 25 pixels and second-nearest-neighbor distances ≥ 15 pixels **(Fig. S2C)**. Minifilament length was defined as the Euclidean distance between the centroids of the paired spots.

To validate the accuracy of this pipeline, we analyzed GATTA-SIM 140B nanorulers (GATTAquant GmbH, Braunschweig, Germany). The same pairing strategy was applied, with adjusted thresholds to account for the smaller expected inter-pole distance and higher particle density (nearest-neighbor ≥ 1 pixel; second-nearest-neighbor ≥ 10 pixels; (**Fig. S2E-F**). Data were acquired over 20 frames (10.4 s) using mpretrack parameters: masscut = 40000, minimum intensity = 30000, and featsize = 2.

To quantify spatial organization along the anterior–posterior (AP) axis, the cortical region was subdivided into 7 equal bins based on the x-coordinate of detected minifilaments. For each bin, the relative fraction of small, medium, and large minifilaments was computed by normalizing to the total number of detected minifilaments within that bin. Spatial differences between anterior (bins 1–3) and posterior (bins 5–7) regions were assessed using two-sample statistical tests. Global spatial variation across bins was evaluated using one-way ANOVA followed by appropriate post hoc tests as described below.

### Behavior analyses and Lifetime

Analyses of minifilament behaviors in the FBR159 strain were performed during polarity maintenance in the anterior domain, and in AB at 2-cell stage during interphase and metaphase, allowing stage-specific comparisons of minifilament behavior. Analysis of minifilament behavior in S1 mutants FBR283 and FBR284, and corresponding control FBR103 strain were performed exclusively at the 2-cell interphase, as lower cortical density facilitates reliable single-minifilament tracking.

Data were collected during the first ~150 s of each stage, corresponding to the period with the highest frequency of minifilament recruitment. To minimize sampling bias, manual selection of minifilaments was performed every 100 frames (~52 s), which approximates the 95th percentile of cortical myosin lifetime.

A minimum separation distance of 15 pixels (~600 nm) between minifilaments was required for inclusion in the analysis, ensuring that trajectories from neighboring structures were adequately assigned. Minifilament selection and trajectory analysis were performed using a custom semi-automated Fiji macro (NIH, Bethesda, MD, USA). The manual component consisted of identifying each minifilament at its first appearance at the cortex and marking its last detectable frame before dissociation into the cytoplasm. The script then extracted minicrops centered on each minifilament, within which behaviors were manually annotated. For each annotated event, the macro automatically quantified net displacement (end-to-end distance) and duration at pixel-scale resolution; for rotational behaviors, angular changes were measured instead of linear displacement.

Quantitative analyses were performed on datasets acquired at a temporal resolution of Δt = 520 ms, which was selected to limit photobleaching while preserving sufficient temporal resolution for behavioral classification. To assess the impact of temporal sampling, additional acquisitions were performed at higher temporal resolution (Δt = 120 ms). Although these conditions were not used for systematic quantification due to increased photobleaching, all behavioral categories were consistently observed, and no substantial differences in minifilament lifetime or event duration were detected. These controls support the robustness of the reported dynamics across imaging conditions.

### Description of computational model of actomyosin networks mechanics

In this study, we employed a well-established agent-based model to simulate actomyosin networks, based on the Langevin equation (J. Li et al., 2017; Jung et al., 2015; T. Kim et al., 2009; Mak et al., 2016). It simulates actin filaments (F-actin), motors, and actin-crosslinking proteins (ACPs) as cylindrical segments. Their spatial positions (x, y, z) evolve according to Brownian dynamics via the Langevin equation. This includes deterministic forces such as bending and extensional forces to maintain structural integrity, as well as repulsive forces to account for volume exclusion between F-actins, preventing spatial overlaps.

F-actin dynamics begin with nucleation, followed by polymerization at the barbed end and depolymerization at the pointed end. ACPs bind to F-actin at a constant rate, with unbinding governed by force-dependent kinetics based on Bell’s law. Motor proteins bind to F-actin and move toward the barbed end, with unbinding rates determined by the parallel cluster model (Erdmann et al., 2013; Erdmann and Schwarz, 2012).

Simulations were conducted in a thin computational domain (5 μm × 5 μm × 0.1 μm) with periodic boundary conditions in the x and y directions. In the z-direction, repulsive forces prevent elements from crossing the boundaries. Each simulation begins with the self-assembly of a thin actin network from F-actin and ACPs.

A full description of the model and the access to its code is provided in (Costache et al. 2022).

### Pulse–chase analysis of Dendra2::NMY-2 photoconversion

Embryos were photoconverted at the pronuclear meeting stage (polarity maintenance phase), a developmental time point immediately preceding global cortical minifilament unbinding, during which minifilaments transiently detach from the cortex and diffuse into the cytoplasm before being recruited again during cytokinetic ring formation.

Prior to pulse–chase experiments, we validated the efficiency of Dendra2 performing a full-field cortical photoconversion. Embryos were imaged in TIRF mode and simultaneously acquired in both green and red channels to monitor the conversion of myosin minifilaments. Photoconversion was induced using a 405 nm laser at 100% power for 3 s.

For pulse–chase experiments, embryos were mounted on high-precision coverslips cleaned by overnight incubation in 1 M HCl, followed by extensive rinsing in absolute ethanol. A defined region of interest (ROI) at the embryo cortex was selected by partially closing the field diaphragm in the illumination path, generating a circular illumination area. Photoconversion within this ROI was induced using a 405 nm laser at 100% power for 3 s of continuous illumination.

### Spots detection and tracking of minifilament intensity after photoconversion

For Fig. 6, to reduce shot noise, we averaged each movie over groups of five consecutive frames, and we subtracted background subsequently. The background image was estimated as the average of 50–100 frames selected during the clearing phase (stage where minifilaments undergo global redistribution into the cytoplasm). We then subtracted the resulting average background image was from the entire movie. To localize spot positions, we processed each movie using the Kilfoil lab algorithm (Crocker & Grier, 1996). By Kilfoil algorithm we also tracked the spots. To compute spot intensities, we used a custom-made MATLAB function that calculates the intensity of circular patches centered at the localized coordinates. The radius of each patch is automatically adapted based on the local signal intensity.

For each localization (x,y), the local mean intensity is first estimated in a small 3×3 window. A corresponding patch radius is then assigned through a log-based interpolation between reference intensity–radius values, constrained between a minimum radius of 2 pixels and a maximum radius of 10 12 pixels. We then applied a A circular mask, avoiding overlap with previously assigned patches, and computed the total intensity within the patch by summing the pixel intensities. Finally, we subtracted the residual background contribution (background intensity per pixel multiplied by the number of pixels in the patch). We measured the residual background intensity per pixel as the average value measured in a region within the embryo that does not contain detectable spots.

### False positive reduction during measurement of minifilament intensity after photoconversion

For Fig. 6, to minimize false positives arising from coverslip impurities or immobile minifilaments originating from dissection, we applied post-detection filters to the spot traces. Single-frame detections were excluded to remove noise. Because true minifilaments display mobility, either small displacements due to cortical thermal diffusion during maintenance or larger movements during furrow formation, we removed traces with total displacement *d=abs(p*_*i*_*-p*_*f*_*)<0*.*5 pixels*, where *p*_*i*_ and *p*_*f*_ denote the initial and final (x,y) position of the spot. We also excluded highly persistent traces since myosins minifilaments turnover on a time scale of ~20s. Specifically, traces detected after frame 50 were removed if present for more than 50 frames (32.5 s), and traces detected within the first 50 frames were removed if lasting more than 70 frames. An upper intensity threshold was further applied, as false positives were often abnormally bright. From frame 50 onward, an intensity cut-off of 7000 a.u. was used. During the first 50 frames—when photoactivated patches reach intensities up to ~10^5^ a.u., a linearly decreasing threshold was applied: *I*_*cut-off*_=15×10^5^ – 2850 x frame.

Additionally, during the first 50 frames, only traces located within a manually defined ROI encompassing the photoactivated region were retained. Manual validation on five randomly selected frames from four movies showed that the initial false positive rate (29%) was reduced to 6% after filtering, increasing the proportion of validated true spots from 71% to 94%. Approximately 5.5% of true spots were excluded by these criteria **(Fig. S5E)**.

### Kinetic model for minifilaments signal after photoconversion

To interpret the temporal evolution of the fluorescence intensity in photoactivated embryos, we developed a simple kinetic model describing the number of fluorescent molecules on the cortex (*N*_*f*_) as function of the binding ( *k*_*on*_), the size of fluorescent cytoplasmic reservoir ( *R*_*f*_(Fig. 6G)) and the unbinding rate (*k*_*off*_). Fluorescent molecules located on the cortex progressively dilute due to exchange with the cytoplasm, where non-fluorescent molecules are present (*R*_*n*_). As time progresses, these non-fluorescent molecules can bind to the cortex, forming a cortical non-fluorescent population (*N*_*n*_). Within a reaction-limited framework, this behavior can be described by the following first-order system of equations:

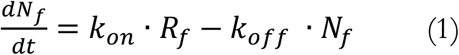

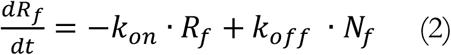

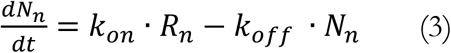

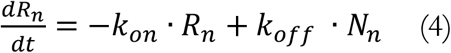

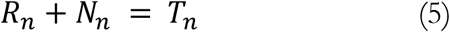

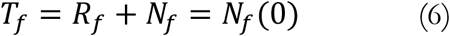

Equations 1 and 3 describe the time evolution of fluorescent /non-fluoresct molecules on the cortex. Equations (2-4) describe the corresponding dynamics of the fluorescent/non-fluorescent molecules in the cytoplasm. Equations (5-6) express conservation of the total number of molecules: (*T*_*f*_) and non-fluorescent (*T*_*n*_) stay constant for the time window considered. remains constant within the time window considered. In particular, the total number of fluorescent molecules is equal to the initial number of fluorescent molecules present on the cortex, (*N*_*f*_ (0)).

The system of Equations (1-6) can be solved analytically obtaining at the equilibrium:

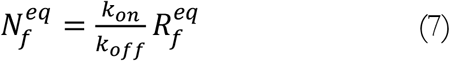

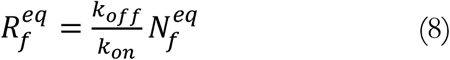

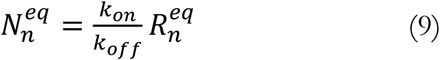

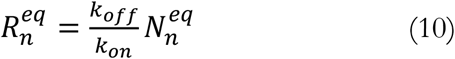

The fraction of fluorescent molecules *I*(*t*) that we show in Fig. 6B is:

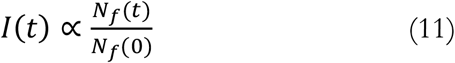

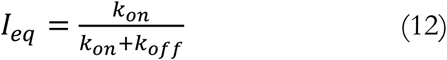

We interpreted *I*_*eq*_as proportional to the plateau intensity observed in **Fig. 6B** after the dilution decay. The true plateau is lower because the experiment only images a portion of the cortex. Based on the geometry of the embryo and the imaging setup, we estimate that roughly one-third of the cortical surface is visible. Therefore, the measured plateau intensity corresponds to only a fraction of the total cortical signal, and we approximate *I*_*eq*_~3 × *I*_*plateau*_.

From super-resolution data, we estimated an average lifetime of the minifilament ranging between 20–25 s. We can then estimate the off-rate (k_off_) of motors from the cortex as *k*_*off*_ = 1/*lifetime* = 0.044 *s*^−1^.

To estimate the cytosolic reservoir R, we refer to the equilibrium relationship 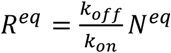. We can reasonably estimate *N*^*eq*^ considering that it is proportional to the estimated number of minifilaments photoactivated in the initial ROI. We estimate the proportionality factor as *α* = *A*_*ph*_/*A*_*tot*_, where *A*_*ph*_ is the area of photoactivated ROI and *A*_*tot*_ is the total visible area of the embryo. Moreover, we consider that the visible portion of the embryo cortex is about ~1/3 of the total cortical surface.

Therefore, we can write *N*^*eq*^~3 × *α* × *N* (0), and consequently 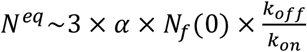. Knowing *k*_*off*_, we estimate *k*_*on*_, from the value of the plateau that follows the decay of the fraction of minifilaments **(Fig. 5A)**, resulting in *k*_*on*_ = 0.019 *s*^−1^. Let us assume that the minifilaments remain intact, without dissolution into monomers during cortical internalization. In this hypothesis, fluorescent minifilaments would coexist in the cytosolic pool with non-fluorescent minifilaments, with a fraction *ff*: 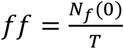 with *T* = *N*^*eq*^ + *R*^*eq*^, therefore 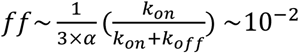. At the furrow, we expect to observe 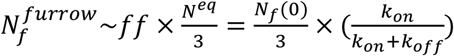 If we estimate the fraction of visible spots relative to the initial photoactivated spots, denoted 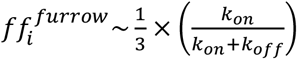 and using 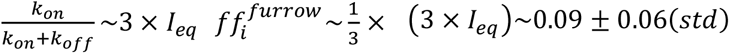, where *I*_*eq*_ is the plateau after the initial dilution of the fluorescent signal (Fig. 6C).

### Determination of the NMY-2 monomer number per minifilaments

Quantification was performed at the anterior pole and in the AB cell during polarity maintenance, 2-cell interphase, and 2-cell metaphase, specifically in regions exhibiting low minifilament crowding to minimize signal overlap and intensity overestimation. We first parameterized the analysis during Polarity maintenance and subsequently applied the same analytical framework to 2-cell interphase and 2-cell metaphase. Polarity maintenance was selected as the calibration stage because it represents a temporally stable phase in which cortical minifilament density remains relatively constant for an extended period. This stability reduces fluctuations in filament number and organization, thereby improving the accuracy and reproducibility of intensity-based quantification and providing a robust reference for comparative measurements across stages.

Images to determination of the number of NMY-2 monomers per minifilaments were collected on TIRF as previously. Briefly, embryos expressing NMY-2 fused with eGFP from the endogenous *nmy-2* locus LP162 (*nmy-2(cp13[nmy-2::eGFP + LoxP]) I*, Dickinson *et al*, 2013) were imaged for short movies (20 frames, 50ms per frame), ensuring that each NMY-2 myosin heavy chain incorporated in the minifilaments were bearing a functional eGFP molecule. We next imaged embryos from cyk-1(jme14[cyk-1::eGFP]) III, that were heavily photobleached, ensuring that CYK-1::eGFP molecules were individual eGFP molecules (Robin *et al*, 2014). We estimated spot intensities by fitting each localization with a two-dimensional Gaussian model including a constant background term. The integrated intensity of each spot was computed as I = 2πAσ^2^, where A is the fitted amplitude and σ the Gaussian width. To avoid artifacts, we excluded spots located closer than a predefined exclusion radius (5 pixels) from another localization, as well as spots whose fitting region extended beyond the image boundaries. In addition, we retained only localizations with fitted σ below a threshold proportional to the typical point spread function width (σ cutoff of 1.8 pixels ≈ 1.5× the median σ) to remove poorly fitted, overlapping, or defocused spots. We applied the same fitting and filtering criteria to both formins and myosin datasets.

We obtained the monomer intensity calibration by computing the mode of intensities of the formins spots distribution **(Fig. 2C)**. Finally, we expressed the intensity of spots in the myosin sample in equivalent monomer units by dividing their fitted intensity by this mode value **(Fig. 2D)**. To assess the robustness of the intensity estimation with respect to the fitting window, we repeated the Gaussian fits using circular fitting regions with radii ranging from 3 to 7 pixels. The estimated number of monomer spots reached a plateau for radii ≥5 pixels; therefore, we selected r = 5 pixels as the fitting radius for the final analysis.

Single-molecule intensity measurements are sensitive and can be subject to variations depending on minute changes in the optical path, changes in laser output, and other difficult-ot-control variations. To avoid experimental drift between the two experiments, instead of direct computation of minifilament intensity, we used an alternative method. Simply put, we used the previously-measured median number of motor domains (n = 30) obtained during maintenance phase at the polarity maintenance as a standard, comparing the intrinsic intensities at all other considered developmental stages (2-cell interphase, 2-cell metaphase) to compute the number of minifilaments.

### Statistical Analysis

Statistical analyses were performed in MATLAB. Normality was assessed using the Shapiro-Wilk test, and homogeneity of variances was evaluated with Levene’s test. Based on these assumption tests: For normally distributed data with equal variances, one-way ANOVA was performed followed by Tukey’s HSD post hoc test. For normally distributed data with unequal variances, Welch’s ANOVA was used followed by Games-Howell post hoc test. For non-normally distributed data (regardless of variance equality), the Kruskal-Wallis test was applied followed by Dunn’s multiple comparisons with Bonferroni correction. Significance was defined as ns, P > 0.05; *P < 0.05; **P < 0.01; ***P < 0.001.

## Acknowledgements

We are grateful to Anna Carvalho for providing the worm strains. We thank the IBPS imaging facility, particularly France Lam and Chloé Chaumeton, for their invaluable technical support in TIRF microscopy. We thank the members of the Robin and Labouesse laboratories for insightful discussions and technical assistance. We are particularly grateful to Simon Begnaud for his contribution to the development of data analysis tools. This research was supported by Pronabec scholarships for Peruvian students under the Bicentennial Generation program (awarded to KLCE), as well as by the ATIP/Avenir and ERC-Tremplin programs (awarded to FBR).

## Author contributions

FBR conceived the project. SKS generated most of the strains and acquired data for Fig. S1 and Fig. 1 (E-G). KLCE performed all experiments and data analyses. CNP provided molecular reagents used in the study. FBR and TF developed Matlab codes used in this study. TF and TK developed simulations and models. PM and EB provided training and support for all acquisitions on the SIM-TIRF microscope.

