## Supplementary Materials for "Persistent and recyclable myosin II minifilaments drive cortical contractility in the early *C. elegans* embryo"

This document includes:

**Supplementary Figure 1**: Spatiotemporal distribution of myosin subunits during early embryonic stages in *C. elegans*.
**Supplementary Figure 2**: Live SIM-TIRF super-resolution imaging resolves the bipolar engagement and the stereotypical length of myosin minifilaments by subunit labeling.
**Supplementary Figure 3**: NMY-2 minifilaments display coordinated behaviors.
**Supplementary Figure 4**: Quantitative characterization of myosin minifilament behaviors revealed by super-resolution microscopy.
**Supplementary Figure 5**: Photoconversion pulse–chase analysis reveals that myosin minifilaments persist and are not disassembled upon cortical detachment/unbinding.
**Supplementary Table 1**: List of *C. elegans* strains used in this study.
**Supplementary Table 2**: List of oligos used in this study.
**Supplementary Table 3**: List of sgRNA targeting sequences used in this study.
**Supplementary Table 4**: List of plasmids used in this study.

#### Supplementary Figure 1

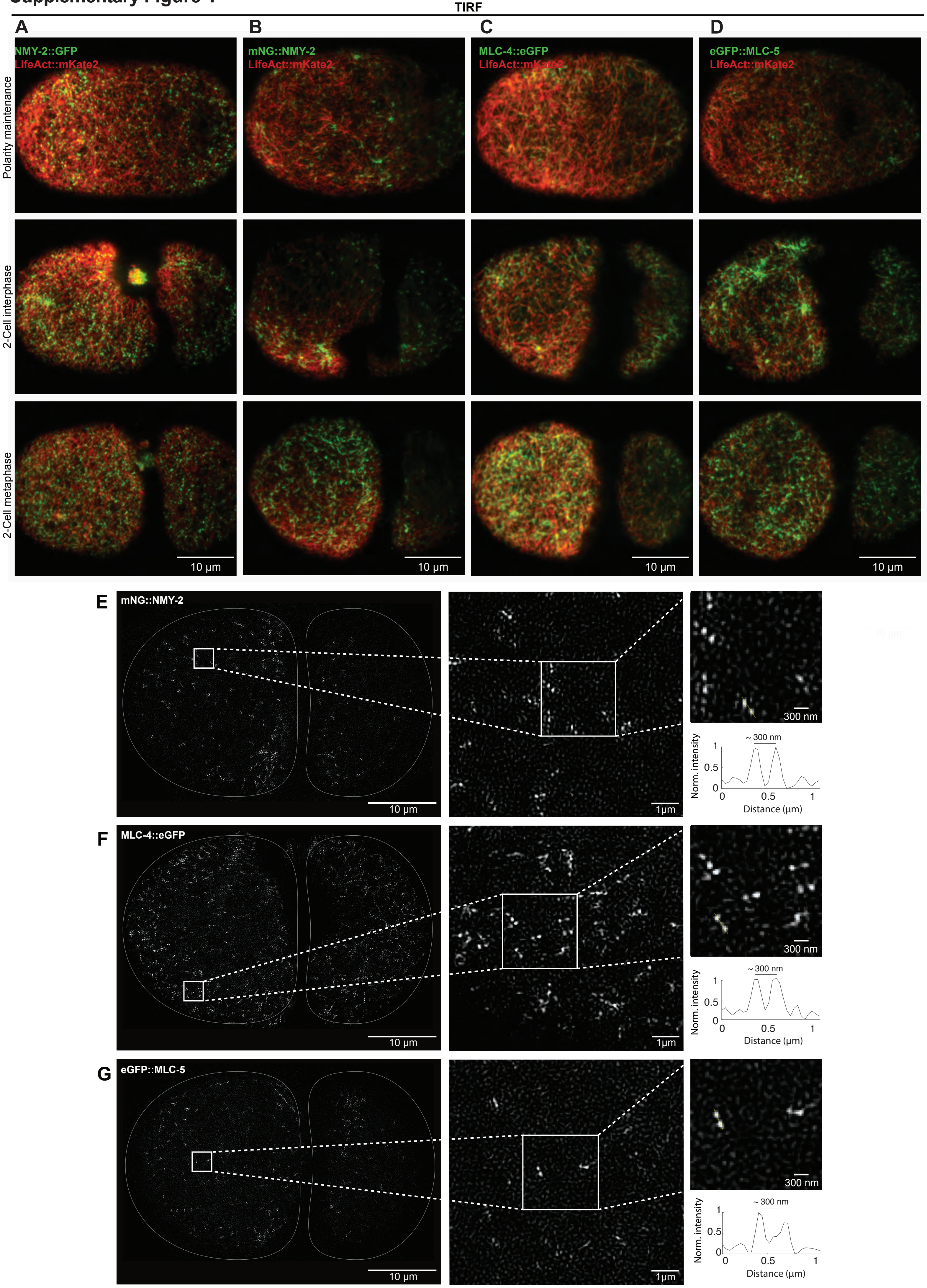

**Supplementary Figure 1. Spatiotemporal distribution of myosin subunits (NMY-2, MLC-4 and MLC-5) during early embryonic stages in *C. elegans*.** Low-magnification TIRF imaging showing the cortical organization and dynamics of myosin subunits during polarity maintenance, 2-cell interphase, and 2-cell metaphase. LifeAct::mKate2 (in red) was used as genetic background. **(A)** C-terminus fusion of myosin heavy chain with GFP, NMY-2::GFP, **(B)** N-terminus fusion of myosin heavy chain with mNeonGreen, mNG::NMY- 2, **(C)** Myosin regulatory light chain fused with eGFP, MLC-4::eGFP. **(D)** TIRF imaging of myosin essential light chain fused with GFP, eGFP::MLC-5. Across all conditions, myosin subunits display a consistent cortical distribution pattern throughout developmental stages. These overview images provide spatial and temporal context for the characterization of individual minifilaments shown in Figure 1. **(E–G)** SIM-TIRF super-resolution microscopy of fluorescent myosin strains. **(E)** mNG::NMY-2, **(F)** MLC-4::eGFP, **(G)** GFP::MLC-5. Individual bipolar myosin minifilaments appeared as *double-dot* patterns of consistent length. Line-scan intensity profiles along the yellow lines drawn across individual minifilaments in higher magnification image revealing an average distance of approximately 300 nm between each pole.

#### Supplementary Figure 2

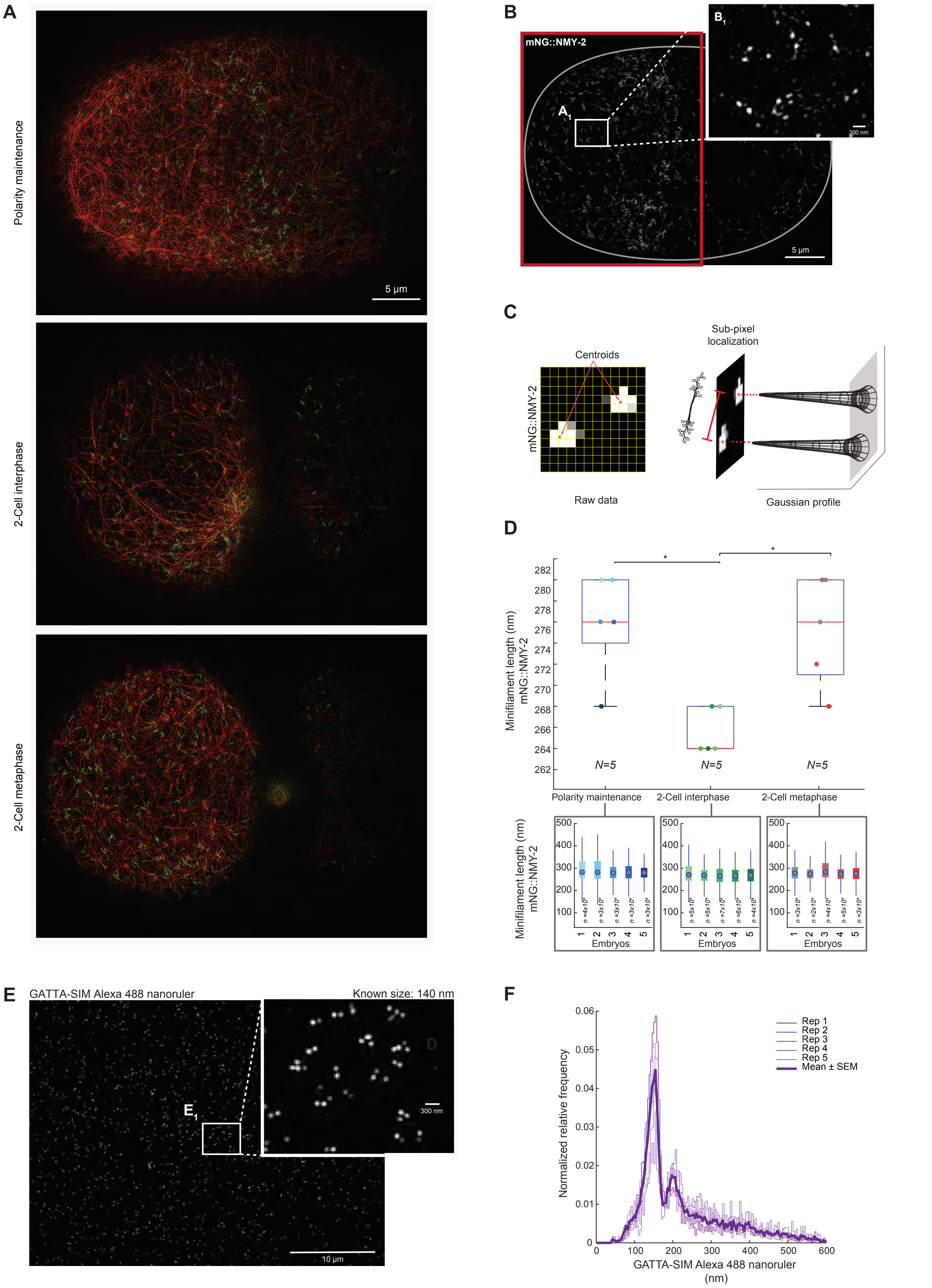

**Supplementary Figure 2. Live SIM-TIRF super-resolution imaging resolves the bipolar engagement and the stereotypical length of myosin minifilaments by subunit labeling (NMY-2, MLC-4, and MLC-5). (A)** SIM-TIRF imaging of N-terminus fusion of myosin heavy chain with mNeonGreen, mNG::NMY-2 resolves individual bipolar minifilaments into two spatially separated poles (*double-dot* pattern). **(B)** Representative embryo during polarity maintenance showing tracked doublets. The red box shows the anterior pole, the area of analysis for this stage **(B1)** Inset shows representative examples of paired minifilament poles. **(C)** Schematic of the procedure for centroid localization. **(D)** Top: Boxplots representing the distribution of per-embryo modal inter-pole distances. N (embryos) = 5 per developmental stage. 2-cell interphase (~264 nm; IQR 264–268 nm), polarity maintenance (~276 nm; IQR 274–280 nm), and 2-cell metaphase (~271–280 nm; *P < 0.05). Individual embryo modal values are shown as colored dots. Modal inter-pole distances were calculated from the full distance distribution using histogram binning (bin width = 0.1 pixels). Bottom: Boxplots of minifilament length per embryo showing comparable dispersion. Each boxplot represents one embryo. n (measurements) > $2 \times 10^4$ for each N (embryos) = 5 embryo per stage. Statistical analyses were performed as described in Methods. Significance levels: ns, P > 0.05; *P < 0.05; **P < 0.01; ***P < 0.001. **(E)** A representative field of view is shown with detected doublets based on measured inter-pole distances. **(E1)** Inset shows representative examples of tracked nanoruler doublets. **(F)** Normalized frequency distribution of nanoruler distances. Individual repetitions are represented by distinct purple tones. The dark solid line indicates the mean normalized distribution, and the shaded regions represents the standard error of the mean (SEM), illustrating variability across repetitions. Histograms show normalized frequency (counts divided by total counts per repetition; bin width = 0.1 pixels). N(repetitions) = 5, n (measurements) > $3 \times 10^3$. Using our MATLAB-based analysis pipeline, nanoruler measurements displayed a bimodal distribution with peaks at 152 nm and ~200 nm, consistent with the manufacturer's validation (yielding 144 ± 6 nm and a secondary peak near 200 nm).

#### Supplementary Figure 3

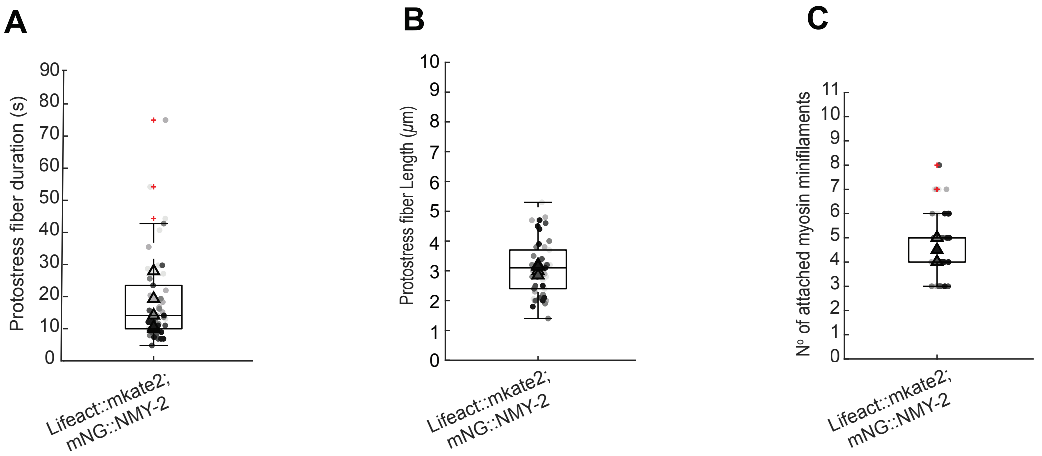

**Supplementary Figure 3. NMY-2 minifilaments display coordinated behaviors. (A)** Bar plots showing the characteristic duration of protostress fibers. **(B)** Boxplots and beeswarm showing the characteristic length of protostress fibers. **(C)** Bar plots showing the characteristic number of attached myosin minifilamemts in protostress fibers.

### Supplementary Figure 4

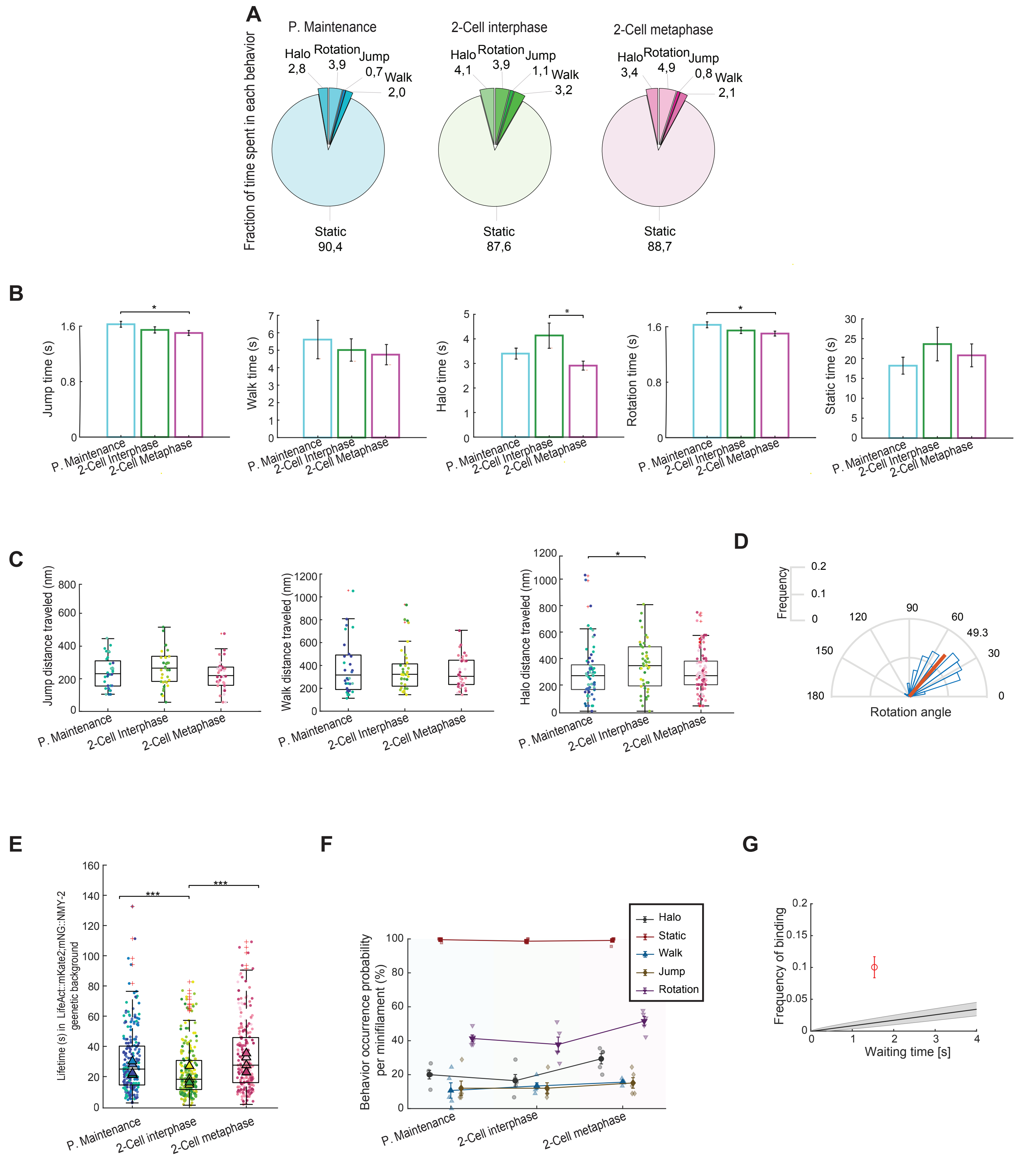

**Supplementary Figure 4. Quantitative characterization of myosin minifilament behaviors revealed by super-resolution microscopy. (A)** Pie charts of time fractions per behavior, showing the distribution of minifilament behaviors across polarity maintenance, 2-cell interphase, and 2-cell metaphase. **(B)** Bar plots showing the characteristic duration of *Jump*, *Walk*, *Halo*, *Rotation*, and *Static* behaviors across polarity maintenance, 2-cell interphase, and 2-cell metaphase. Error bars represent ± 95% confidence intervals (CI). **(C)** Boxplots showing the characteristic distances traveled during *Jump*, *Walk*, and *Halo* behaviors. **(D)** Distribution of the characteristic rotation angle. **(E)** Boxplots showing measured lifetimes (seconds) during polarity maintenance (IQR 14.4- 40.2 s), 2-cell interphase ~18 s (IQR 11.4-30.7 s), and 2-cell metaphase ~28 s (IQR 15.9-45.8 s). N(embryos) = 5 per stage, n(minifilaments/embryo) = 45. **(F)** Probability that an individual minifilament exhibits at least one occurrence of a given behavior during its cortical lifetime. Percentages do not sum to 100 %, as individual minifilaments may exhibit multiple behaviors during a single lifetime. **(G)** Binding frequency of *Jump* compared with the random binding frequency as function of the waiting times.

### Supplementary Figure 5

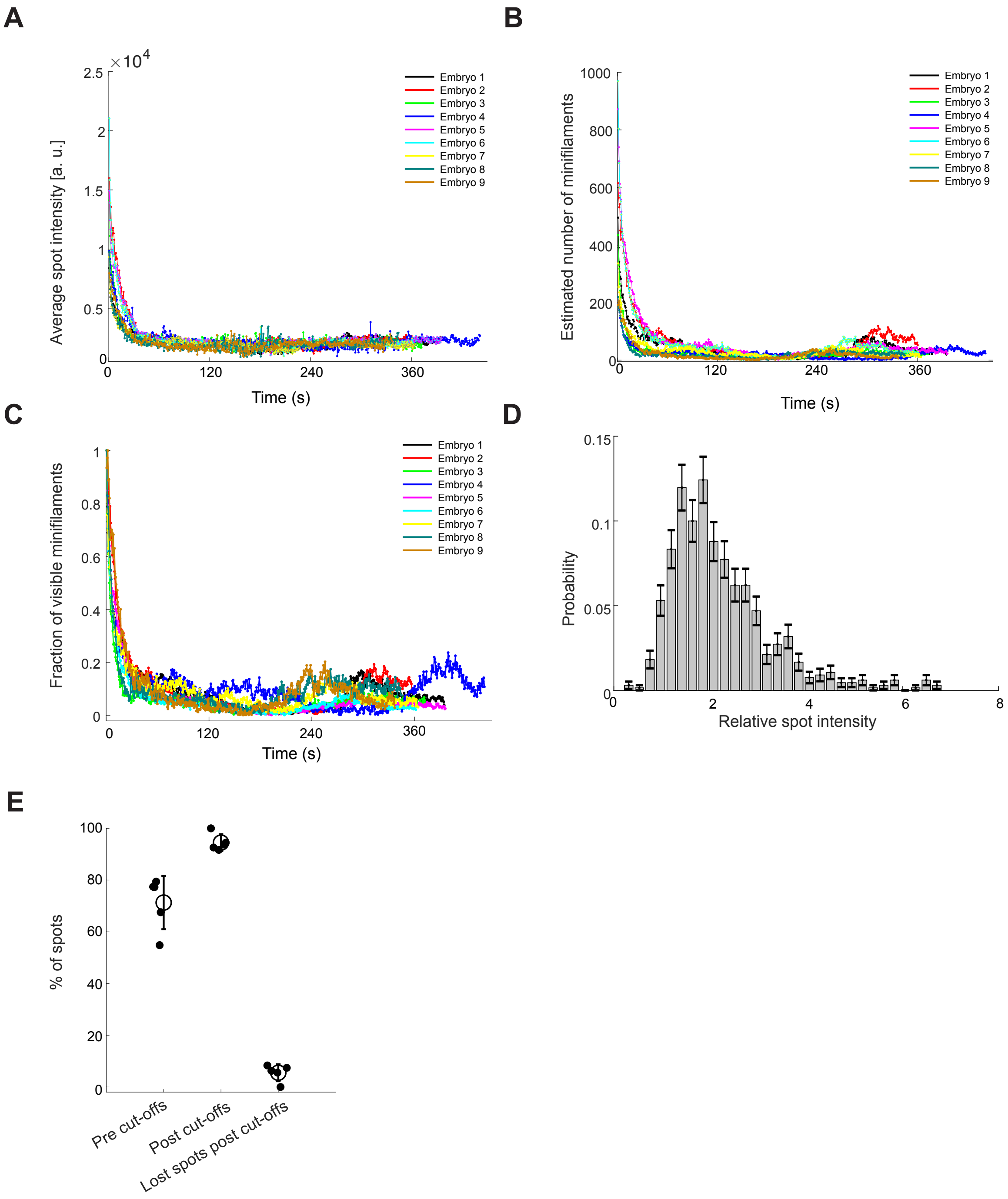

**Supplementary Figure 5. Photoconversion pulse–chase analysis reveals that myosin minifilaments persist and are not disassembled upon cortical detachment/unbinding. (A)** Time evolution of the average spot intensity following photoconversion, from polarity maintenance (pronuclear meeting) to the first cell division. N(embryos) = 9. The initially high signal following photoconversion rapidly decreases within the first ~40 s and then reaches a stable plateau intensity $I_{eq}$ until the end of the movie. **(B)** Estimated number of minifilaments per frame, calculated as the total measured intensity $I$ divided by the plateau intensity value $< I_{eq} >_{time>60s}$, where $< I_{eq} >_{time>60s}$ is the time average of the intensity stating from the first minute till the end of each movie. **(C)** Fraction of visible spots estimated from photoconversion till the first cell division. N(embryos) = 9. **(D)** Probability distribution of spot intensities at 45 s and 60 s after photoconversion (see Fig. 6F), pooling data from all 9 embryos. **(E)** Percentage of manually validated spots before cut-off application, followed by the percentage of validated spots after cut-off application, the residual false positive contamination, and the percentage of true spots excluded by the applied filters. The total amount of spots considered for this evaluation is 186.

**Table S1. List of *C. elegans* strains used in this study.**

| Strain name | Genotype | Source |
|---|---|---|
| N2 | Wild-type Bristol strain | CGC |
| FBR78 | *mlc-4(jme02[mlc-4::mCherry+loxP]) III* | CRISPR/Cas9 |
| FBR96 | *mlc-4(jme04[mlc-4::eGFP+loxP])III* | CRISPR/Cas9 |
| FBR103 | *gesIs001[Pmex-5::LifeAct::mKate2::nmy-2(3'UTR); unc-119(+)]; mlc-4(jme04[mlc-4::eGFP+loxP])III* | Genetic cross: SWG062 x FBR96 |
| FBR119 | *nmy-2(cp52[nmy-2::mKate2+LoxP unc-119(+) LoxP])I; unc-119(ed3)III; mlc-4(jme04[mlc-4::eGFP+loxP])III* | Genetic cross: LP229 x FBR96 |
| FBR140 | *mlc-5(jme10[eGFP^3xFlag::LoxP::mlc-5])III* | CRISPR/Cas9 |
| FBR154 | *nmy-2(cp52[nmy-2::mKate2+LoxP unc-119(+) LoxP]) I; unc-119(ed3)III; mlc-5(jme10[eGFP^3xFlag::LoxP::mlc-5])III* | Genetic cross: FBR140 x LP229 |
| FBR159 | *gesIs001[Pmex-5::LifeAct::mKate2::nmy-2(3'UTR); unc-119(+)]; nmy-2(cp141[mNG^3xFlag::nmy-2])I* | Genetic cross: FBR174 x LP389 |
| FBR160 | *cyk-1(jme14[cyk-1::eGFP]) III* | CRISPR/Cas9 |
| FBR174 | *gesIs001[Pmex-5::LifeAct::mKate2::nmy-2(3'UTR); unc-119(+)]* | Outcrossed from SWG6062 |
| FBR184 | *gesIs001[Pmex-5::LifeAct::mKate::nmy-2UTR, unc-119+]; mlc-5(jme10[eGFP^3xFlag::LoxP::mlc-5])III* | Genetic cross : SG062 x FBR140 |
| FBR207 | *nmy-1(jme17[mNG::nmy-1])X* | CRISPR/Cas9 |
| FBR209 | *nmy-1(jme17[mNG::nmy-1])X; nmy-2(cp52[nmy-2::mKate2 + LoxP unc-119(+) LoxP])I; unc-119 (ed3)III* | Genetic cross: FBR207 x LP229 |
| FBR282 | *nmy-2(Dendra2::nmy-2)I* | SunyBiotech |
| FBR283 | *nmy-2[prt37(R718C)]I;gesIs001[Pmex-5::LifeAct::mKate:: nmy-2(3'UTR), unc-119(+)]; mlc-4(jme04[mlc-4::eGFP+loxP])III* | Genetic cross: GCP375 x FBR103 |
| FBR284 | *nmy-2[prt38(S250A)]I;gesIs001[Pmex-5::LifeAct::mKate:: nmy-2(3'UTR), unc-119(+)]; mlc-4(jme04[mlc-4::eGFP+loxP])III* | Genetic cross: GCP376 x FBR103 |
| GCP375 | *nmy-2[prt37(R718C)]I* | Osório et al. 2019 |
| GCP376 | *nmy-2[prt38(S250A)]I* | Osório et al. 2019 |
| LP162 | *nmy-2(cp13[nmy-2::GFP+LoxP])I* | Dickinson et al, 2013 |
| LP229 | *nmy-2(cp52[nmy-2::mKate2+LoxP unc-119(+) LoxP])I; unc-119(ed3)III* | Dickinson et al, 2017 |
| LP389 | *nmy-2(cp141[mNG^3xFlag::nmy-2])I* | Dickinson et al Genetics 2015 |
| LP572 | *nmy-2(cp278[GFP-C1^3xFlag::nmy-2])I* | Heppert et al 2016 |
| SWG007 | *nmy-2(cp8[nmy-2::GFP+unc-119(+)])I; unc-119(ed3) III; gesIs001[Pmex-5::LifeAct::mKate2:: nmy-2(3'UTR); unc-119(+)]* | Reymann et al., 2016 |
| SWG062 | *gesIs001[Pmex-5::LifeAct::mKate2::nmy-2(3'UTR), unc-119+]; cyk-1::gfp65c* | Reymann et al., 2016 |

**Table S2: List of oligos used in this study.**

| Name | Sequence(5'-3')* | Description | Purpose |
|---|---|---|---|
| OCP-82 | ACAAGGATGAGGCTTAAAC**GTTTAAGAGCTATGCTGGAAAC** | mlc-4 sgRNA forward | Construct sgRNA on U6 vector (pML2840) |
| OCP-83 | GTTTAAGCCTCATCCTTGT**CAAACATTTAGATTTGCAATTC** | mlc-4 sgRNA reverse | |
| OCP-78 | ACGTTGTAAAACGACGGCCAGTCGCCGGCA**CGCTTCCCTTGGAAAAGAAGTCAC** | mlc-4 5' arm forward | Construction of mlc-4::GFP repair template in pJJR82 vector or mlc-4::mCherry repair template in pJJR83 vector |
| OCP-79 | CATCGATGCTCCTGAGGCTCCCGATGCTCC**AGCCTCATCCTTGTCCTTGGTTC** | mlc-4 5' arm reverse | |
| OCP-80 | GATTACAAGGATGACGATGACAAGAGA**TAAACTGGCTCGCAAACTCACAC** | mlc-4 3' arm forward | |
| OCP-81 | CAGGAAACAGCTATGACCATGTTATCGATTTC**AACAGCACACAAGAAACATGGCA** | mlc-4 3' arm reverse | |
| oSUMAN-505 | CTGTACCTGCAGGTACTGC**GTTTAAGAGCTATGCTGGAAAC** | nmy-1 sgRNA forward | Construct sgRNA on U6 vector (pML2840) |
| oSUMAN-506 | GCAGTACCTGCAGGTACAG**CAAACATTTAGATTTGCAATTC** | nmy-1 sgRNA reverse | |
| oSUMAN-499 | TTCCGGATGGCTCGAGTTTTTCAGCAAGAT**ATTCTCCGCCTTTCCAAGCTC** | nmy-1 5' arm forward | Construction of mNeon::nmy-1 repair template in pJET2.1 vector |
| oSUMAN-500 | **TTGTTAACTTCGTAATATCTGAAAATTGTTTTCAATTTTAATTG** | nmy-1 5' arm reverse | |
| oSUMAN-501 | AACAATTTTCAGATATTACGAAGTTAACAAA**ATGGTGTCGAAGGGAGAAGAGG** | mNeon forward | |
| oSUMAN-502 | CTGTACCTGCAGGTACTGCAGATCACCCATGCCGCTACCACTGGT**CTTGTAGAGTTCATCCATTCCCATCAC** | mNeon reverse | |
| oSUMAN-503 | **ATGGGTGATCTGCAGTACCTGC** (PAM mutation) | nmy-1 3' arm forward | |
| oSUMAN-504 | AGAATATTGTAGGAGATCTTCTAGAAAGAT**CGAGGATGAAGCTCTTTGGGAA** | nmy-1 3' arm reverse | |

*Oligos are listed for the constructs used for CRISPR/Cas9 mediated knock-in transgenesis. For each oligo, the region that anneals to the target is in bold. PAM site mutation is shown in red.

**Table S3: List of sgRNA targeting sequences used in this study.**

| Gene | sgRNA sequence* | Location |
|---|---|---|
| mlc-4 | ACAAGGATGAGGCTTAAAC**TGG** | 3'end in ORF |
| mlc-5 | CTGTGAGAAATGGACGATT**TGG** | 5'end in ORF |
| nmy-1 | CTGTACCTGCAGGTACTGC**AGG** | 5'end in ORF |

**Table S4: List of plasmids used in this study.**

| Plasmid | Characteristic | Source |
|---|---|---|
| pDD162 | Peft-3::Cas9 + Empty sgRNA | (Dickinson et al., 2013) |
| PRF4 | rol-6(su1006)] | (Frøkjær-Jensen et al., 2008) |
| pCFJ90 | myo-2p::mCherry::3'UTR unc-54 | (Frøkjær-Jensen et al., 2008) |
| pJET1.2/blunt | Cloning vector | Thermo Fisher Scientific |
| pJJR83 | mCherry^SEC^3xFlag | Addgene |
| dg397 | slot3 Entry mNG::3xFLAG::unc-54 (3'UTR) | (Hostettler et al., 2017) |
| pML2840 | derived from PIK198 | (Suman et al., 2019) |

*PAM is underlined and bold.
